# The gene *ivory:mir-193* controls scale type differentiation in *Heliconius* butterflies

**DOI:** 10.64898/2026.02.24.707831

**Authors:** Joseph J. Hanly, Leo Camino, Julia C. Holder, Paola Calderon-Oñate, Pushkar R. Wagh, Joshue Ruiz, Leia Zhao, Eva D. Stroh, Luca Livraghi, Ling S. Loh, Carlos F. Arias, Lawrence E. Gilbert, Gregory A. Wray, Arnaud Martin, W. Owen McMillan

## Abstract

The *ivory:mir-193* locus is a genetic hotspot underlying melanic wing pattern variation across Lepidoptera, acting as a master regulator of melanic scale fate. This includes the genus *Heliconius*, where aposematic mimicry is driven by three scale cell types: light Type I, dark-melanic Type II, and red Type III scales. We tested functions of *ivory:mir-193* in *Heliconius* using CRISPR-induced somatic mosaic knockouts of *ivory* and *mir-193* across multiple pattern morphs. Knockouts converted Type II scales to Type I scales. Effects on Type III scales varied among and within individuals, indicating that *ivory:mir-193* is permissive for Type III development. Using single-nucleus RNA-seq, we profiled the transcriptional landscape of *H. melpomene*-Δ78k mutants lacking *mir-193*. Loss of *mir-193* produced readthrough transcription, consistent with a model where miR-193 acts as a co-transcriptional terminator. These data show that *ivory* drives wing pattern variation, while *mir-193* mediates downstream diversity in scale fate across Lepidoptera.

## Introduction

Many butterfly and moth species exhibit wing pattern polymorphisms that underlie a wide range of ecological adaptations, including crypsis, mimicry, thermoregulation, and sexual selection ^1–3^. Interestingly, a growing number of melanic polymorphisms have been mapped to a homologous genomic region called *ivory:mir-193* (previously referred to as the *cortex* locus), in a wide range of species, including in aposematic mimicry polymorphism in *Heliconius*^4,5^, the peppered moth industrial melanism polymorphism^6^, cryptic leaf-masquerade in *Kallima* butterflies^7^, regulation of plasticity in buckeyes^8^, and many others^9–11^. This unprecedented scale and level of replication highlight the *ivory:mir-193* microRNA locus as an exemplar genetic hotspot^12,13^ where colour polymorphisms are repeatedly driven by allelic variation across Lepidoptera. In all described cases, *ivory:mir-193* is involved in the regulation of colour in melanic scales but not of other pigments^14–17^. As the association of a miRNA locus to adaptive phenotypes sheds new light on the mechanisms behind evolutionary change, and because of its striking repeatability across at least 110 MY of divergence, it is fundamental to gain insights into how this locus regulates wing colouration phenotypes.

*Heliconius* is a genus of neotropical butterflies that includes 48 species with many cases of within-species pattern divergence and between-species convergence. Almost all the variation in *Heliconius* wing patterns can be attributed to the spatial arrangement of just three scale cell types, rather than to a continuum of colour states: the white or yellow “Type I” scales; the heavily melanised “Type II” scales; and the “Type III” scales that contain red and orange-brown ommochrome pigments^18–20^. Importantly, each scale type is characterised by a combination of pigment composition, intracellular organisation and cuticular ultrastructure, reflecting developmentally specified cellular identities^21,5^ (**Fig. 1A-B**). The spatial arrangement of these three scale types generates bold, high-contrast patterns that function in aposematic signalling, increasing conspicuousness against the forest background and promoting predator learning and avoidance.^22,23^. Mapping of wing pattern variation in *Heliconius* has identified a small toolbox of genes responsible for the spatial arrangement and specification of these scale cell types (**Fig. 1C**), including the transcription factor genes *optix*, which specifies red Type III scales^24,25^; and *aristaless-1* (*al1*) which blocks the uptake of the yellow pigment 3-OHK and acts as a switch between yellow and white Type I scales^26^.

**Figure 1:**
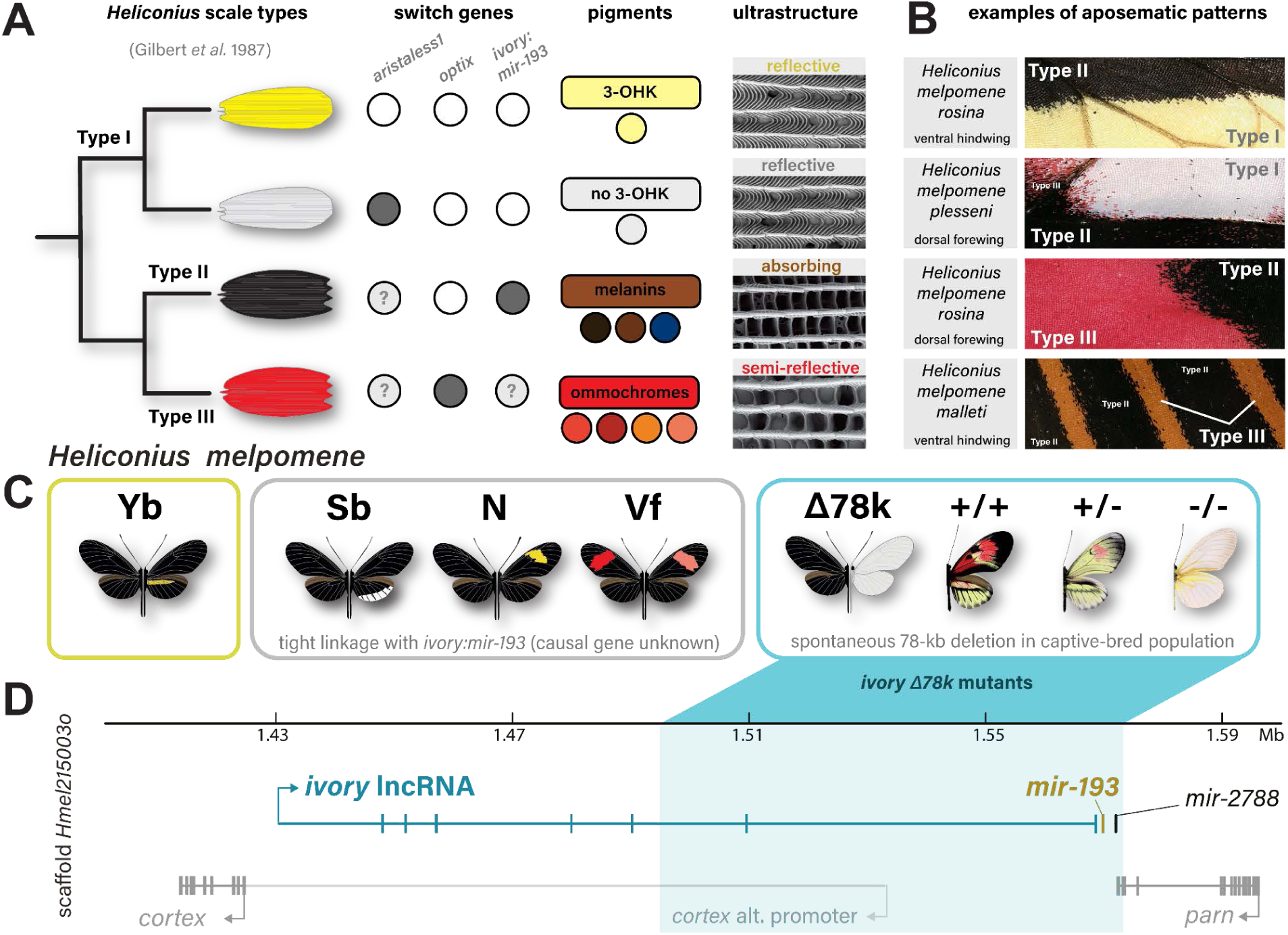
Scale type specification in *Heliconius* butterflies. (**A**) Scale cell type tree, as initially conceptualised by Gilbert et al., annotated with switch genes for cell types identified by^15,24,26^, and showing a sample range of colours for specific scale types depending on population of origin, as well as scanning electron micrographs of upper scale surfaces, showing nanostructural variation between types. (**B**) Shows examples of how the three scale cell types can be combined in high contrast arrangements to form aposematic pattern elements found in multiple *Heliconius* co-mimics. (**C**) The named pattern alleles identified by linkage mapping, all of which are tightly linked with the *ivory:mir-193* locus in *H. melpomene*. (**D**) A schematic depiction of the *ivory:miR-193* locus, demarking the 78kb deletion previously identified in an insectary mutant.

Convergent, patterned changes in the distribution of Type I scales have occurred in the co-mimics *H. melpomene* and *H. erato,* as exemplified by the presence or absence of the hindwing yellow bar (**Fig. 1C**). In both species, this variation was mapped to a locus containing the protein-coding gene *cortex*^4,5^. In addition, an insectary mutant line showing a near-complete conversion of Type II/III to Type I scales mapped to a 78 kb deletion (hereafter *Δ78k*) spanning the same genomic region^14^ (**Fig. 1C-D**). Subsequent re-annotation of the locus revealed the presence of the long non-coding RNA *ivory*, which runs antiparallel to *cortex* and overlaps with its 5’UTR. Similarly to other species, CRISPR/Cas9-mediated somatic knockouts of *ivory* in *H. erato* cause high-penetrance transformation of melanic Type II scales to yellow-white Type I scales across all cuticular surfaces, and expression of *ivory* closely prefigures the presence of Type II scales in pupal wings^15,27^.

Key work in *B. anynana* then elucidated a relationship between *ivory* and *mir-193.* The 3’ end of the *ivory* transcript is adjacent to both *mir-193* and *mir-2788*^28^, and *ivory* acts as their primary transcript^29^. In *B. anynana*, knockout lines of *mir-193*, but not of *mir-2788*, phenocopied *ivory* knockout phenotypes, and both *ivory* and *mir-193* were co-expressed in melanic wing pattern elements. This led to a working model where the *ivory* lncRNA acts as a primary transcript that templates the expression of *mir-193*, with mature *miR-193* then acting as a regulatory factor that modulates melanin production. The role of *mir-193* has not yet been formally tested in *Heliconius*, and two observations hint that the *Bicyclus* model is not fully in agreement with the genetic effects of the *ivory:mir-193* locus in this genus. First, the transformation of Type II (black) scales into Type I (yellow-white) induced by *ivory* KOs can be decomposed into several effects. At the pigment level, *ivory*-deficient crispant tissues not only lose dark melanin but also acquire 3-OHK absorption^15^, implying that regulatory effects go beyond the control of melanin pathway genes. At the ultrastructural level, these scales shift from a light-absorbing, porous surface to a light-reflecting surface, also hinting at regulatory effects on ultrastructural components of scale development. In addition, *ivory* KOs and the *Δ78k* deletion also affect red Type III scales^14^. Together, these preliminary data suggest that *miR-193* acts as a trans-regulatory factor modulating cell type specification rather than just melanin production.

In this paper, we sought to directly test the versatile functions of *ivory:mir-193* in *Heliconius* butterflies through the production of mKOs of both *ivory* and *mir-193* in multiple species and pattern forms, with particular attention to the fate of red Type III scales. Additionally we examined transcriptional effects of this locus by performing single nucleus RNA sequencing on individuals from the Piano Keys line with different allelic states of the *Δ78k* deletion, allowing us to evaluate the *trans*-effects of *ivory:mir-193* in *Heliconius*, and to assess conserved functional roles of this element with other species.

## Results

### Knockouts of *ivory* and *mir-193* cause loss of dark scales

To examine the relationship between *ivory* and *miR-193* in *Heliconius*, we generated crispants for both the first exon of *ivory* and *mir-193* seed site (Table S1), in multiple pattern forms of *H. melpomene, H. erato* and *H. numata*. Crispants for both *ivory* and *mir-193* displayed clones with complete conversion of melanic Type II scales to yellow or white Type I scales (**Fig. 2**). The conversion of Type II to yellow or white Type I depended on pattern morph. For example, *H. m. plesseni* forewings displayed Type II to Type I yellow conversions exclusively, whereas in *H. m. rosina* forewings, scale type conversion was position-dependent, with proximal scales converting to yellow and distal scales to white.

**Figure 2:**
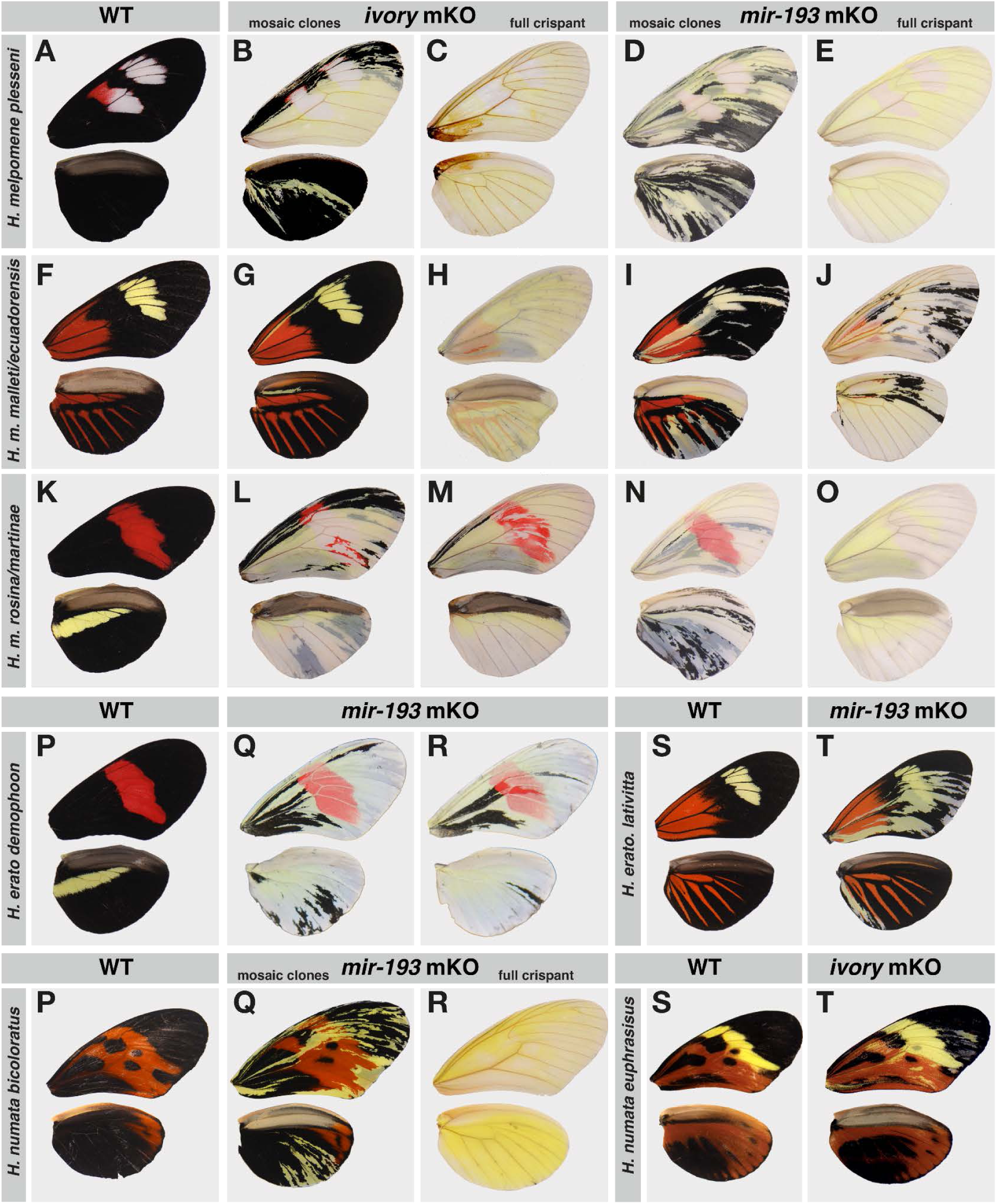
Phenotypic effects of *ivory* and *mir-193* mKO in *Heliconius* butterflies. Ventral wild-type wings, plus multiple *ivory* and *mir-193* mKO phenotypes for *H. melpomene plesseni* (A-E), *H. m. malleti/ecuadorensis* (F-J), and *H. m. rosina/martinae* (K-O). mKOs of *mir-193* in *H. erato demophoon* phenocopy previously published knockouts of *ivory* (P-R) plus *mir-193* knockout in *H. e lativitta*.

Additionally, we knocked out *mir-193* in other nymphalid species where we had previously knocked out *ivory*, additionally confirming the phenocopy of *ivory* (**Fig 3**). In *H. erato, Junonia coenia* and *Vanessa cardui*, *mir-193* crispants also closely mirror those of *ivory*, causing conversion of melanic scales into light coloured scales. Notably, in *V. cardui*, as well as multiple other species^15^, the knockout of *ivory:mir-193* solely affects melanic scales and does not have any apparent effect on ommochrome-containing red or orange scales, whereas in both *J. coenia* and *Heliconius spp*., the red Type III scales are also perturbed.

**Figure 3:**
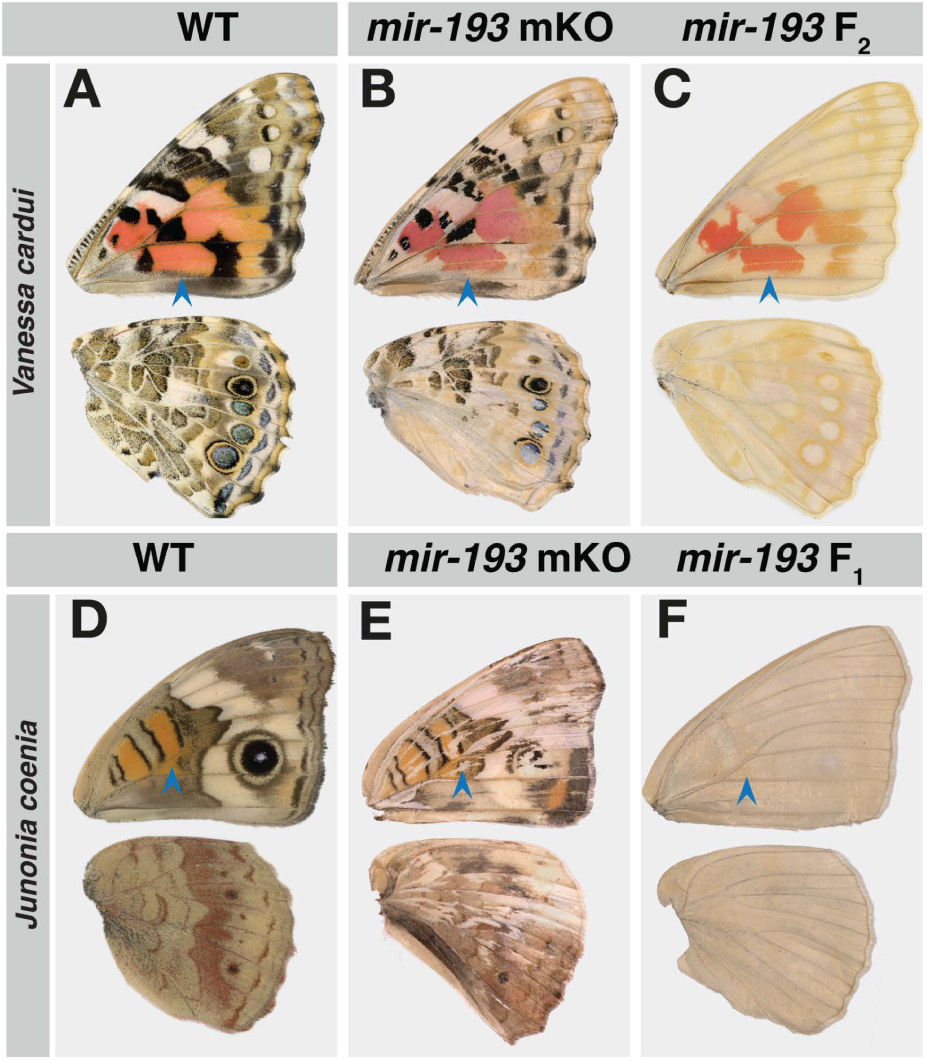
*mir-193* knockout in additional nymphalid species phenocopies the knockout of *ivory*. -A-B show ventral wild-type and *mir-193* mKO in *Vanessa cardui*, while C shows F_2_ homozygous mutant. Blue arrowheads indicate an ommochrome-containing red pattern element which retains red pigmentation in both mKOs and germline mutants. D-E show *Junonia coenia* ventral wild-type and F_1_ *mir-193* mKO, with blue arrowheads indicating a ommochrome-containing red element - in E, a clone that intersects through this element loses red pigmentation, while in F, all red pattern elements lack red pigment.

The effects of *ivory* and *mir-193* knockout on Type III scales were much more variable than the effects on Type II scales, falling into four main classes: we either observed 1) Type III scales becoming pale pink, 2) pink Type III scales accompanied by large punctate clumps of red pigment, 3) Type III scales becoming folded along the long axis, causing a deformed, taco-like shape, 4) complete conversion of Type III scales to Type I scales. When the transformation from Type III to Type I is complete, most pattern forms exhibited a shift to white, except for in *H. m. malleti* where they became yellow. In some individuals, we saw clones of more than one class on the same wing surface, indicating that neighbouring mutant clones can exhibit differing expressivity of the mutant phenotype, with one clone retaining some ommochrome pigment and the other displaying complete conversion to Type I (**Fig. 4**). This suggests that these neighbouring clones could contain different CRISPR-induced deletions that induce differing phenotypes, likely corresponding to true loss-of-functions and hypomorphs of different penetrance.

**Figure 4:**
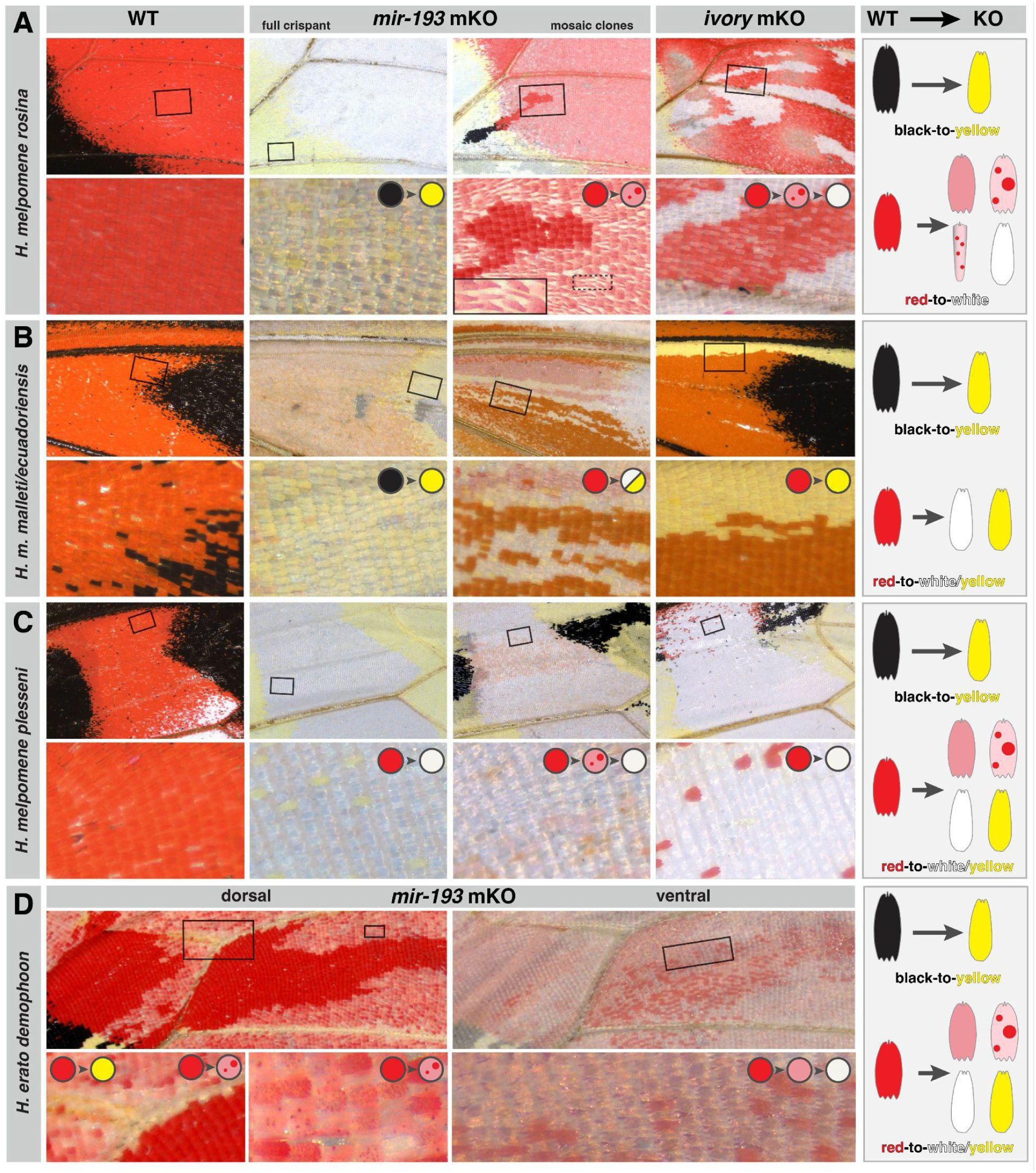
*ivory* and *miR-193* are required for differentiation of black Type II scales and permissive for the normal differentiation of red Type III scales. (**A**) High magnification images of the forewing red band region in *H. melpomene rosina*. The top row shows spatially matched regions at 10x, adjacent to for a wild-type, *mir-193* crispants, and an *ivory* crispant. The black inset rectangles are shown at 50x magnification in the lower panels. Inset coloured circles show the transition of scale type from wild type to mutant in the pictured region, and the rightmost panel shows all observed transitions, including Type II to Type I, and Type III one of several malformed Type III-like scales or to Type I. (**B**) depicts the same range of images for *H. m. malleti/ecuadorensis*, while (**C**) shows *H. m. plesseni.* (**D**) shows high magnification images of a *mir-193* crispant *H. erato demophoon*.

Wing crispant clones cannot be directly genotyped as the scale cells degenerate towards the end of pupal development^30^. As such, we genotyped individuals with different extents of clonality from thoracic tissue, and observed proportions of coverage reduction that correlated with the extent of the loss of Type II scales. (**Fig S5, Table S3**).

### Single-nucleus RNA-seq of *mir-193* mutants

Our CRISPR results illustrate that *ivory:mir-193* function can be context dependent in different pattern forms, and produce several classes of scale transformations. Therefore, in order to examine the transcriptomic consequences of mutations at this locus on scale type specification, in a context where all scales were transformed to Type I, we opted to use the previously described “Piano Keys” line of *H. melpomene*, which carries a recessive, 78-kb deletion that removes the 3’ portion of the *ivory* lncRNA and the whole of *mir-193*. Homozygous mutant *Δ78k* butterflies phenocopy the most extensive knockout phenotypes of *ivory:mir-193*, with all Type II and the vast majority of Type III scales converted to Type I scales.

Pupal wings were dissected, snap-frozen and genotyped by PCR, allowing the selection of individuals who were either *Δ78k* -/- or +/+ for snRNA-seq analysis. We selected one at 30% of pupal development wild-type ‘Piano Keys’, and three *Δ78k* homozygotes at 31%, 41% and 55%, and combined this with a previously-generated time series of wild-caught *H. melpomene* collected in Panama, in which clusters had been identified and validated^31^. The full analysis includes 11,743 nuclei sequenced to an average depth of 56,250 reads per nucleus.

Following integration and normalisation of samples, we used unsupervised clustering and annotated cell types using our previously identified and validated markers for pupal wing tissues^31^. As the phenotype of *ivory* and *mir-193* KO specifically affects wing scales, we focused our analysis on the Sense Organ (SO) lineage (**Fig. 5A-D**)^31,32^. This lineage includes SO precursors (SOPs), which express marker genes *delta, sns* and *N-cad*, and which divide at around 20% development into scale-forming cells expressing the markers *sv and ss,* and socket-forming cells expressing markers *Sox15* and *Su(H)*. Epithelial cells expressing *E-cad* and *Notch* (**Fig. S6**), as well the marginal cells expressing *fz3* and *cut,* were excluded from the analysis.

**Figure 5:**
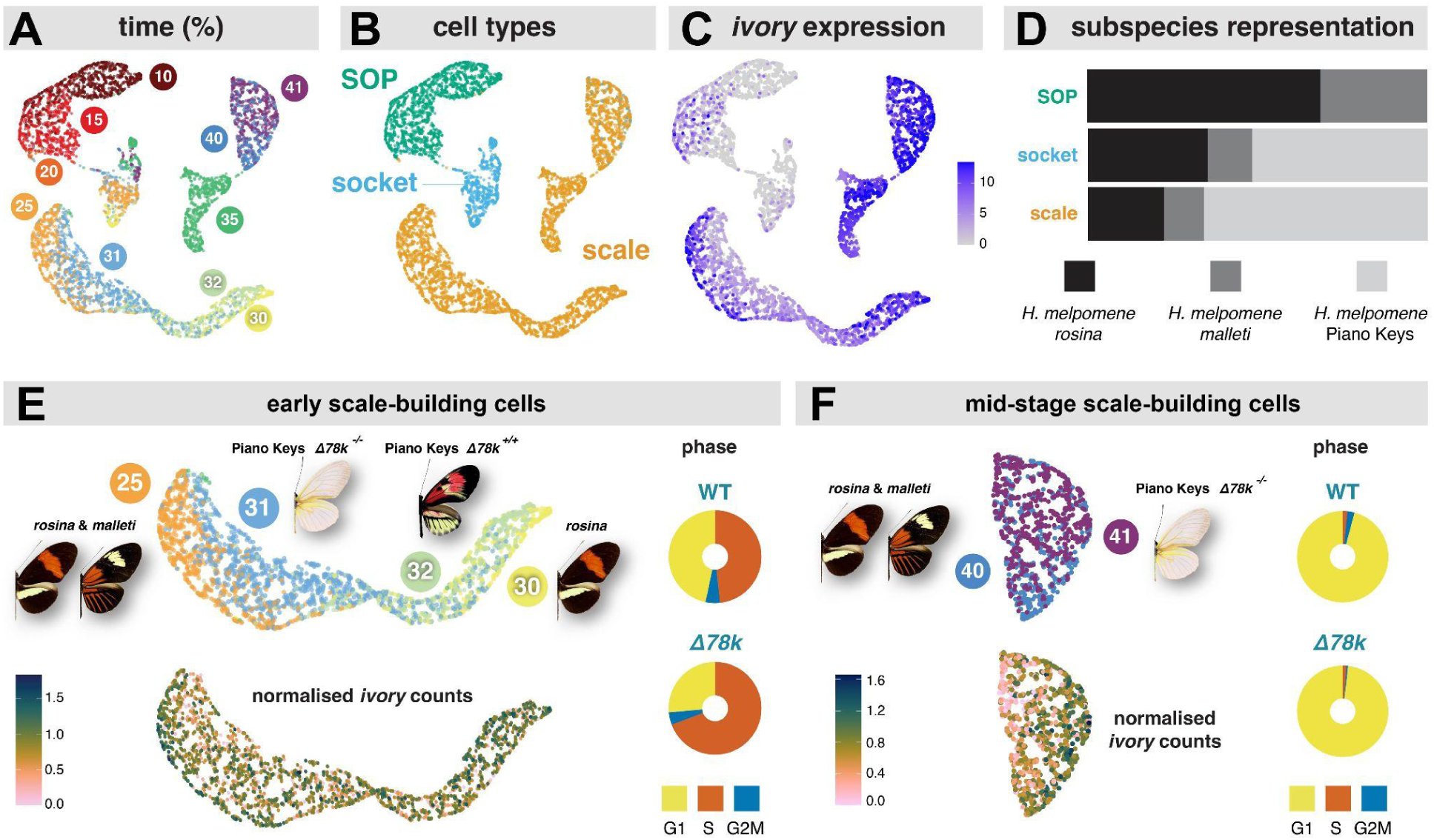
snRNA-seq of developing wild-type and mutant wings. (**A**) depicts time series of SOP-lineage cells from wings of wild types and mutants. (**B**) shows cell type assignment based on markers described and validated by Loh et al (2025). (**C**) shows expression of ivory throughout the time series. (**D**) the distribution of cell types between integrated sample groups; SOP cells are only present in 10% and 15% samples, and so are represented by only the wild-type *H. m. rosina* and *malleti*, whereas socket and scales were recovered in roughly equal proportions between the wild-type and PK samples. (**E and F**) highlight the early scale-building cell cluster (E) and the late scale-building cell cluster (F). The upper panels indicate the phenotype at percentage pupal development of the individuals making up each cluster; the lower panel indicates per-sample normalised ivory expression, used for defining the high vs low ivory groups. To the right are donut charts indicating the proportions of cell cycle phase of the wild-type cells vs mutant cells.

Expression of *ivory* initiates stochastically in SOP cells, and becomes very high in scale-building cells while remaining low in socket-building cells (**Fig. 5C**). Scale-building cells from the Piano Keys individuals fell into two groups. The first, ‘early’ group contains all scale-building cells from wild-caught *H. melpomene* from 25% and 30%, plus all scale-building cells from the Piano Keys wild-type 30% individual, and all scale-building cells from the Piano Keys *Δ78k* 31% individual (**Fig. 5E**. The second later group contains scale-building cells from the 40% wild-type plus the 41% mutant (**Fig. 5F**). No scale-building cells were recovered from the 55% *Δ78k* mutant sample, and so it was dropped from further analysis.

We previously described how cell cycle phase imputation of scales indicates that early post-specification scale building cells go through a period of S-phase with no progression to G_2_M-phase, which corresponds to the increase in ploidy of scale cell nuclei^31^. Here, we note that a higher-than-expected fraction of our scale precursor nuclei from mutant individuals were in S-phase, which may indicate differences in the regulation of cell cycle progression in the *mir-193*-null state (**Fig. 5E**). At the later time point, almost all scale-building cells in both wild types and mutants are in G_1_, indicating that they have likely exited the endoreplicative cell cycle (**Fig. 5F**).

### Transcriptional dynamics at the *ivory* locus

We examined per-cell read alignments at the *ivory* locus (**Fig. 6A, Fig. S6**). A combination of high expression, high intronic read accumulation, and the long introns of *ivory,* leads to *ivory* being the second most abundant transcript in the scale precursor nuclear transcriptome (after the mRNA binding protein *heph*) (**Fig S7**). This stands in stark contrast to previous observations from bulk tissue RNA-seq from *H. melpomene* and other nymphalid species, where *ivory* was observed at very low abundance^15^, or was completely undetectable in RNA-seq on wild-type tissues despite being detectable by qPCR^29^.

**Figure 6:**
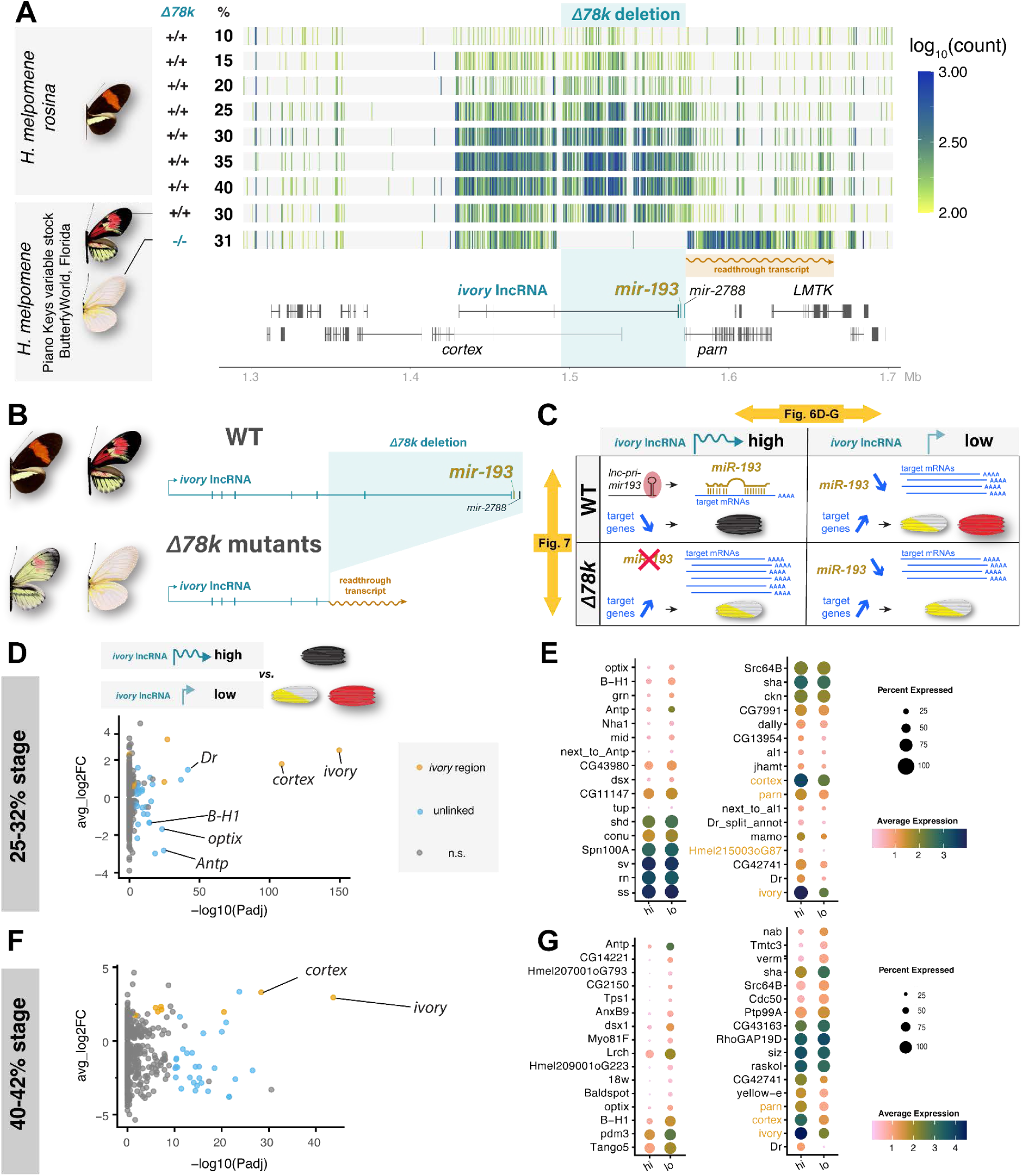
*ivory:mir-193* locus expression dynamics in single nucleus RNA-seq data. (**A**) ribbon plots for each individual snRNA-seq sample at the *ivory:mir-193* locus, showing only scale-building cells, and only the depth of reads aligning in the forward orientation (*i.e.* in frame with the *ivory* transcript). In the wild-type individuals we observed a gradual accumulation of reads across the exons and introns of *ivory* over time. In both the heterozygous and mutant individual, no reads mapped within the deletion, but high levels of reads mapped in the forward orientation after the end of the deletion and terminating at the gene *LMTK*, which we refer to as the ‘readthrough transcript’. (**B**) depicts the *ivory* and *ivory*-readthrough transcripts. (**C**) A schematic depiction of what happens in a wild-type “high *ivory*” vs “low *ivory*” state (upper panels) vs. what happens in each state in a *Δ78k* mutant scenario where *mir-193* is absent. (**D, F**) Volcano plots of differential expression between early scale forming cells (D) and mid-development scale-forming cells (F), with high *ivory* vs. low *ivory*, regardless of genotype, identifies the differential expression of several transcription factors with known roles in scale development. (**E, G**) Dot plots highlighting the top differentially expressed genes; several of the same genes are differentially expressed at both time points.

**Figure 7:**
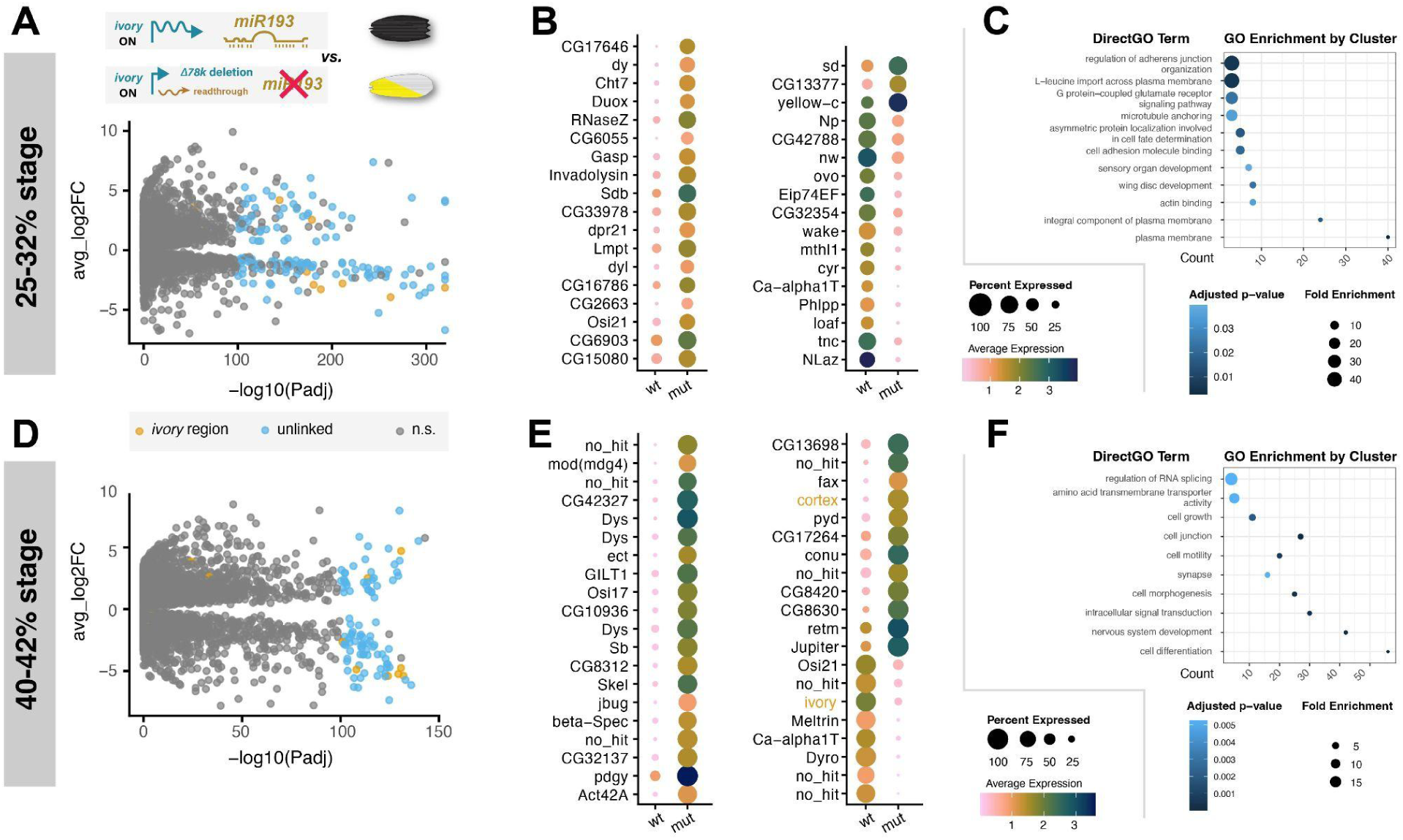
Differential expression between wild-type and *mir-193* mutant scales with high *ivory* expression. **(A-C)** Volcano plot of differential expression in early scale-building cells, dotplots highlighting the top differentially expressed genes, and GO term analysis results. (**D-F**) Volcano plot, dotplot and GO term plots for mid-development scale-building cells.

Notably, a large number of intronic reads are observed for this transcript in scale-building cells— pre-mRNA reads are expected to accumulate in single-nucleus RNA sequencing due to the relative enrichment of nuclear components, in comparison to whole-cell or bulk RNA sequencing where processed mRNAs in the cytoplasm are expected to dominate^33^. Accumulation of reads mapping to *ivory* exons and introns initiated at the 10% stage (**Fig. 6A**), and in wild-type butterflies, we observed read accumulation over time.

The *Δ78k* deletion does not encompass the promoter of *ivory*, but does include much of the 3’ region, including *mir-193* and *mir-2788*, and in read mapping in *Δ78k* snRNA samples, we observed that transcription initiation of *ivory* is not impeded by the deletion, but the resultant transcript is aberrant. In wild-type wings, read accumulation ends around *mir-193*, and just 3’ of the gene *parn*. In contrast, the *Δ78k* mutants exhibit high, continuous intronic and exonic read accumulation that begins at the *ivory* promoter, and continues after the deletion breakpoint at high levels up to the gene *LMTK*, at which point the intronic read accumulation ends (**Fig. 6A-B**). This matches the aberrant transcript described in *B. anynana* bulk RNA-seq, where the deletion of *mir-193* resulted in failure of transcription termination and the generation of a spliced transcript that also extended to and terminated at *LMTK*^29^.

### Expression differences between scale types

Given that we observed heterogeneous *ivory* expression in the nuclei of both wild type and mutant scale-building cells, we split the data into “high *ivory”* and “low *ivory”* populations. We inferred that the “high *ivory*” population were scale-building cells fated to become Type II scales, while the “low *ivory*” were fated to become either Type I or Type III scales. Cells. We performed pseudobulk differential expression analysis between “high” and “low” populations, thereby identifying factors that are potential upstream activators or repressors of *ivory* and Type II differentiation.

In the early cluster, we identified several factors known to be involved in scale fate specification and colour in *Heliconius* or other systems, including the transcription factors *optix* (which specifies Type III scales), *al1*, *B-H1* and *Antp*, as well as *mamo*, which has been linked to pigment deposition in the larvae of *Bombyx mori*^34^ (Fig 6D-E). In the late cluster, we identified several of the same transcription factors in the same patterns, including *optix, B-H1, Antp, Dr* and *ds*, as well as *pdm3*, a repressor of dark scale fate in *V. cardui* (Fig 6F-G). We also saw differential expression of the pigmentation gene *yellow-e*, which was higher in the high-ivory cells, consistent with an early accumulation of melanin in presumptive Type II scales.

### Expression differences between wild-type and mutant scales

Given that we could detect signals consistent with the differentiation of presumptive Type II scales, we then considered differential expression between wild-type “high *ivory*” cells and *Δ78k* mutant “high *ivory*” scales, thus comparing presumptive Type II cells that would develop correctly, to presumptive Type II cells that lacked *mir-193* and would therefore fail to differentiate (*i.e.* scales that would become Types II/III *vs.* scales that would become Type I). As miRNAs function through the targeted degradation of mRNAs, genes with higher expression in mutants than in wild-types are candidate direct targets of *miR-193*.

We again performed pseudobulk differential expression. Several of the top DE genes are immediately 3’ of *ivory*, likely reflecting the long aberrant transcript that terminates at *LMTK*. Notable overexpressed genes include the *osiris* family of endomembrane-associating genes, the yellow homolog *yellow-c*, and the transcription factor *ovo/svb*. We examined functional enrichment in the over-expressed gene set using PANGEA for GO Term analysis, and ModPhea^35^, a database of phenotypes observed upon gene knockout in *Drosophila* as recorded on FlyBase. This gene set was enriched for terms for Actin binding, microtubule binding, cell adhesion molecule binding and adherens junctions. The knockout phenotype enrichment included genes that affect development of ‘inner photoreceptor cells’ and ‘scutellar bristles’ (supplementary data). In contrast, the set of genes that is reduced in expression in “high ivory” *Δ78k* mutants is enriched for GO Terms related to steroid hormones, Toll signalling, and wing morphogenesis, while the knockout phenotype enrichment includes terms for ‘death before end of second instar’ and ‘anterior wing margin’.

## Discussion

We identified extensive parallels in the mechanism of the *ivory:mir-193* locus between *Heliconius* butterflies and other nymphalids, providing concurring evidence that: *(a) ivory* is a transcriptional hotspot locus driving natural variation in wing pattern; *(b) mir-193* is required for the termination of the *ivory:mir-193* primary transcript; *(c)* the mature miRNA *miR-193* is the *trans*-acting factor generated by this locus that modulates several aspects of scale differentiation. It is highly likely that this mechanism is conserved between the many Lepidoptera in which this locus has been mapped as a master locus of melanic variation. Additionally, we highlighted that beyond a simple involvement in melanin pathway synthesis, this transcriptional unit is required for normal scale type specification including black and red scale types.

### Conserved transcriptional dynamics

Our data corroborates a working model where *ivory* lncRNA acts as a noncoding precursor transcript (lnc-pri-miRNA) that templates the expression of *mir-193*, as described in *Bicyclus*^29^, and is in concordance with the observation a substantial fraction of miRNAs are derived from within an lncRNA host gene both in humans^36^ and in other insects^37^. The promoter region of *ivory* is highly conserved^15^, and likely interacts with different *cis-*regulatory modules to initiate the patterned expression of the *ivory* transcript, ultimately causing accumulation of mature *miR-193* which then acts as a downstream *trans*-acting factor causing scale cell type differentiation. Interestingly, snRNA-seq revealed that *ivory* is the most abundant transcript in scale cell nuclei at the stages we assessed, hinting that an early and robust activation of this locus is occurring well before the onset of scale pigmentation.

Once initiated, how is the elongation of the *ivory:mir-193* nascent transcript terminated? In the canonical mode of miRNA biogenesis described in mammalian cells, miRNAs are produced from lncRNA primary transcripts that, rather than stopping shortly after a polyadenylation signal^38^, are co-transcriptionally terminated at the site of miRNA hairpin excision^39,40^. Consistent with this model, our snRNA-seq data indicates that the *mir-193* region is required for the termination of its primary transcript: in *Δ78k*^-/-^mutants, the absence of *mir-193* results in read-through transcription, with RNApolII continuing to the next transcriptional terminator in the same orientation on the chromosome, which is *LMTK*. Similarly, in short *mir-193* knockouts in *B. anynana*, a long, spliced transcript was observed that initiated at the conserved *ivory* promoter and extended to *LMTK*, though in wild-type transcriptomes, no *ivory* transcript was observed, potentially because its level of expression was too low to be detected^29^. These similarities, along with the high sequence conservation of the *ivory* promoter sequence and the conserved synteny in this locus, suggest that it could be a conserved mechanism in Nymphalidae and possibly throughout Lepidoptera.

The observed ectopic accumulation of *ivory* transcript when *mir-193* is absent implies that the *ivory* lncRNA alone is unlikely to have a molecular function beyond acting as the lnc-pri-miRNA of *miR-193*, but in spite of this, the *ivory* promoter and first exon is conserved in almost all assayed lepidopteran genomes^15^, raising the possibility that the coupling of *ivory* and *mir-193* is highly conserved, with *ivory* acting as the transcriptional regulator and *mir-193* as the downstream *trans*-acting factor, and that this mechanism evolved early in the diversification of the Lepidoptera.

### ivory:mir-193: necessary for Type II, permissive for Type III scale fate

Uniformly, across three tested species of *Heliconius* and three other nymphalids, targeted deletions of *ivory:mir-193* cause the loss of melanic scales, referred to in *Heliconius* as Type II scales, and this is consistent with *B. anyanana* and the Tea Geometrid *Ectropis grisescens*^41^. In contrast, the effects on ommochrome-containing scales (Type III scales in *Heliconius*) are more variable between lineages. In *V. cardui*, and *B. anynana*, *optix*-responsive, ommochrome-containing red scales are unaffected by *ivory:mir-193* perturbation, whereas in *J. coenia*, ommochrome-containing red regions are fully converted to light-coloured scales^25,27,42^.

In *Heliconius spp.*, the range of possible outcomes for Type III scales we observed could be explained as a spectrum of expressivity, with a range of effects from regular Type III scale gross morphology but a lighter colour; large ‘clump-like’ deposits of ommochrome; a folded ‘taco-like’ appearance; or a complete transformation to Type I scales. This diversity in clone types was observed even within single wings. A likely explanation is the presence of dosage effects—if *ivory* or *miR-193* crispants are hypomorphs rather than complete loss of function mutants, then some miR might still be produced at low levels. If a low level of miR is needed for the proper formation of red scales, then while total loss of the miR causes complete transformation of red scales to yellow or white, a small remaining quantity of miR could permit the formation of either aberrant red scales or even of normal red scales. Supporting this model, Piano Keys *Δ78k* heterozygotes retain aberrant red scales whereas *Δ78k* homozygous mutants show the complete conversion of red scales to white or yellow. Future studies involving fine-tune control of miR levels could assess whether the effect of *miR-193* on downstream mRNA targets is dosage-dependent, and whether this ultimately has threshold effects on proper scale differentiation and pigmentation.

### ivory:mir-193 and the evolution of cell type and phenotype

The different functions of *ivory:mir-193* in different lineages indicates that the set of downstream targets of *miR-193* is likely to evolve through the acquisition or loss of miRNA seed sites in target genes, leading to rewiring of gene regulatory networks responsible for scale cell fate specification^43^. It is likely that there is an ancestral functional involvement in the specification of dark scales, as all of the cases where *ivory:mir-193* has been mapped involve changes to the extent of melanic patterns.

Target sites for some conserved miRNAs have been observed to evolve slowly, and so it is possible that there are a core set of downstream conserved targets^44^, which could in future be analysed through comparative transcriptomics and bioinformatic seed site prediction. In at least the two lineages where we observe an additional role in red ommochrome scales (*Heliconius* and *Junonia*), there may have been a burst of acquisition of new target genes of *miR-193* through the gain of seed sites in 3’UTRs specific to regulation of red scale types^44^. Alternatively, a role in red pigmentation may be ancestral to the nymphalids, accompanied by a pattern of loss in other tested lineages. This will be resolvable by looking at conservation and phylogenetic signal in 3’UTR seed sites, once they are identified.

Within *Heliconius*, the variation between pattern forms that has been mapped to *ivory:mir-193* has different effects in each of the species studied here. In *H. numata*, all pattern variation maps to this locus, including regional effects on Type I, Type II and Type III scales; in *H. melpomene*, the locus causes local switching in pattern between Type I and Type II scales, while also being involved in the spatial arrangement of Type III scales only in the red forewing band; in *H. erato*, variation at *ivory:mir-193* solely affects the specification of Type I and II scales. In spite of that, the perturbation phenotypes of *ivory:mir-193* are remarkably similar between the three species, indicating that we should expect to see little variation within the genus in the downstream targets of *miR-193*. It is therefore likely that the differences in patterning functions in wild populations are driven by upstream regulatory variation.

Notably, there is minimal intersection between differentially expressed genes identified here and in previous bulk RNA-seq studies comparing wild types to *ivory*-null^27^ and *miR-193*-null pupal wings^29^. There are a number of explanatory factors; first, time point selection was done independently and for different purposes, and it is possible the spectrum of target genes shifts over time during wing development. Second, there is likely to be churn, drift and selection for different targets in different butterfly species, and the gain and loss of different sets of targets could be responsible for phenotypic evolution in scales and patterns. This highlights that we might expect to find signatures of selection in the 3’UTRs of coding genes.

## Conclusion

We have shown that *ivory:mir-193* is required for proper differentiation of multiple scale cell types in *Heliconius* wings. More broadly, scales are a good model for studying cell type individuation because of the high diversity in cell types, tractability of the developing cells, and extensive pre-existing genomic and phenotypic data, making this system well placed for the future study of the origin and diversification of novel cell types^45^.

## Acknowledgements

We thank Remi Mauxion, Oscar Paneso, and Daniel Romero for extensive assistance with butterfly collection, stock rearing, and plants in Panama, as well as Emily Breckenridge at UT Austin. We also thank Kiana Kamrava and Alexander Carter for support with CRISPR-Cas9 experiments at GWU. We thank Ron Boender of Butterfly World, FL for generously providing living specimens for study by the Gilbert Lab at UT Austin. Library preparation and sequencing was performed with assistance from Castle Raley at the GW Genomics core, and sequencing performed by the Duke University Sequencing and Genomic Technologies (SGT). We also thank the GWU HPC team for computing infrastructure^46^.

## Methods

### Butterfly husbandry

*Heliconius* butterflies originated from a breeding farm located in Ecuador, except for *H. melpomene rosina* and *H. erato demophoon*, which were collected on Pipeline Road in the vicinity of Gamboa, Panama. Stocks of butterflies were established and maintained at the *Heliconius* Stock Center and Rearing Facility situated at the Smithsonian Tropical

Research Institute in Gamboa, Panama. Adults were kept in large (3m x 3m 3m) outdoor net cages with a rain protection roof. Each cage was provided with nectary flowers for adult feeding, including *Lantana camara, Psychotria poeppigiana,* and *Psiguria triphylla,* and butterflies were also fed daily a mix of sugar water and pollen. Females were given a specific *Passiflora* host plant for oviposition, depending on the species, and caterpillars from these stocks were raised on big *Passiflora* plants until pupation. The rearing temperature ranged between 28-34°C and humidity between 70%-85% depending on the season. The necessary butterfly collection permits were obtained from the Ministry of the Environment of Panama, according to the Panamanian government and STRI regulations.

A captive population of *Heliconius melpomene* f. “Piano Keys” butterflies in which the *Δ78k* mutation segregates was a gift of Ron Boender (Butterfly World, Coconut Creek, Florida), maintained at the UT Austin Department of Integrative Biology greenhouses by Larry Gilbert. The Piano Keys phenotypes resulted from artificial selection on marginal hindwing white pattern elements at Butterfly World circa 1990, and the *Δ78k* mutation spontaneously occurred in this stock around 2013^14^. “Pale Piano Key” mutants (in retrospect, these were heterozygotes for miR-193 KOs) arose in that stock at Butterfly World in 2015, and were soon shared with LEG in Austin. Homozygous “Ivory” segregates soon appeared at BW and later in Austin where investigations by JH, AM and LEG, began in 2019.

*J. coenia* and *V. cardui* colonies were maintained in environmental growth chambers at 28 °C and 25 °C, respectively, under 60–70% relative humidity and a 14:10 h light:dark photoperiod. Egg collection, larval rearing on artificial diet, and microinjection procedures were performed as previously described^47,48^. *V. cardui* larvae were obtained from Carolina Biological Supply, and *J. coenia* larvae were provided by Robert D. Reed from a laboratory colony originally established by Fred H. Nijhout.

### CRISPR reagents

Cas9 single-guide RNAs (sgRNAs) were designed to target the promoter/exon 1 region of the *ivory* lncRNA and the hairpin region of *mir-193*, using reference genomes from LepBase (v4) and NCBI Genomes (*V. cardui*, GCF_905220365.1; *J. coenia*, GCA_018235105.1). Synthetic sgRNAs were obtained from Synthego and Integrated DNA Technologies, resuspended in Low TE buffer (10 mM Tris–HCl, 0.1 mM EDTA, pH 8.0) at 500 ng/μL, aliquoted (2.5 µL), and stored at −70 °C. Recombinant Cas9-2xNLS protein (PNABio) was resuspended at 1,000 ng/μL in *TE* buffer supplemented with 0.05% cell-culture–grade phenol red, aliquoted (2.5 μL), and stored at −70°C. CRISPR injection mixes were prepared immediately prior to use by combining equal volumes of Cas9 and sgRNA aliquots to yield final concentrations of 500 ng/μL Cas9 and 250 ng/μL sgRNA.

### CRISPR microinjections

Syncytial embryos were microinjected within 2 hr after egg laying for all species. *Heliconius* eggs were glued to microscope slides using Elmer’s glue and injected with Cas9:sgRNA duplexes using borosilicate glass capillary needles mounted on a XenoWorks Digital Microinjector (Sutter Instrument). Hatched larvae were kept in individual plastic cups and fed with the appropriate host plant *ad libitum*. The wings of adult crispants were digitized in standardized conditions and stored in glassine envelopes for further analyses.

### PCR and Nanopore-Based Genotyping of CRISPR Mutants

DNA for genotyping was extracted from ethanol preserved thorax tissue. Genomic DNA was isolated using the Qiagen DNeasy Blood & Tissue Kit following the manufacturer’s protocol, and final DNA quality and concentration were assessed using a Nanodrop or a Qubit fluorometer (). PCR genotyping was performed to verify CRISPR induced mutations within the ivory genomic region. Primer pairs were designed to amplify a 732 bp region surrounding the CRISPR target site. Primer designs were generated using either the Heliconius melpomene version 2.5 reference genome or the Heliconius erato demophon v1 genome within Geneious Prime® 2025.2.2, targeting conserved sequences flanking the edited locus (Table S1). PCR reactions were conducted using primer specific annealing temperatures and conditions listed in Table SMX. Amplicons were visualized on 1.5% agarose gels to confirm successful amplification, and positive reactions were used directly for library preparation.

Nanopore sequencing libraries were prepared using the Rapid Barcoding Kit 96 V14 (SQK RBK114.96) and sequenced on a Flongle R10 series flow cell according to Oxford Nanopore Technologies’ standard protocols. Basecalling and demultiplexing were performed with Dorado v1.2 (https://github.com/nanoporetech/dorado) using the Sup high accuracy model on a GPU enabled node of a HPC. Demultiplexed reads were mapped to either the *H. melpomene v2.5* or *H. e. demophon v1* ivory reference scaffold—selected according to the species under analysis—using minimap2 with parameters optimized for ONT long reads. Mapping depth and coverage were calculated using samtools^49^. A custom R script was used to generate coverage profiles and visualize read depth and genomic coordinates across the entire ivory locus.

### Isolation of Δ78k deletion carriers

For single nucleus transcriptomics, individuals from the *H. melpomene* f. “Piano Keys” population, in which the *Δ78k* deletion segregates, were reared in a greenhouse environment through their mid-fifth instar, and moved to a laboratory (22-23°C) in plastic containers and with *Passiflora biflora* cuttings to finish development. Pupation time was recorded and pupae were dissected at stages ranging 25-55%, based on a total pupal developmental time averaging 228 hr at room temperature for non-mutant individuals, and 242-254 hr for mutants. Forewings were snap-frozen in liquid N_2_ and stored at −70°C while the rest of the pupal body was preserved in DESS solution (20% DMSO, 0.25 M EDTA, and saturated NaCl, pH 8.0). A PCR assay was used to genotype individuals for the presence and allelic dosage of the *Δ78k* deletion, using one pair of primers that amplified the distal promoter of *cortex*, within the deletion (pair 1, expected amplicon size 318 bp) and one pair that spanned the deletion itself (pair 2, expected amplicon 121 bp). Individuals were called homozygous wild-type if only pair 1 gave an amplicon of expected size, homozygous mutant if only pair 2 gave an amplicon of expected size, and heterozygous if both primer pairs gave an amplicon of expected size.

### Single-nucleus RNA sequencing

This study includes snRNA-seq samples in 7 previous Heliconius melpomene samples Nuclei preparations were performed as previously published on all samples (Loh et al Development 2025). Briefly, frozen forewings were thawed, lyzed using dounce homogenizers before clean-up by density-based centrifugation in a 1.8 M sucrose solution (Sigma-Aldrich, NUC-201), and counted using Trypan Blue staining on a hemocytometer. Nucleus suspensions were diluted to between 700 and 1,800 nuclei/μl and library preparation was performed with a targeted recovery of 5,000 nuclei. Library construction was carried out using the Chromium Single Cell 3′ Gene Expression v3 reagent kit (10x Genomics). Final cDNA libraries were quantified with Qubit fluorometry, and fragment size distributions were evaluated using the Agilent 2100 BioAnalyzer with the High Sensitivity DNA kit (5067-4626). Pooled libraries were sequenced on an Illumina NovaSeq 6000 S4 flow cell, yielding an average of ∼281 million total reads per sample, corresponding to approximately 56,250 paired-end 100-bp reads per nucleus. Sequencing was performed by the Sequencing and Genomic Technologies (SGT) core facility at Duke University. Single-nucleus raw reads are available on the NCBI repository under BioProjects PRJNA1404329 and PRJNA1118807.

### snRNA-seq data analysis

BCL files were converted using bcl2fastq2 (RRID:SCR_015058) and aligned to the *H. melpomene melpomene* v.2.5 genome^50^ with the *H. melpomene melpomene* v.3.1 annotation^51^. The annotation orthology assignment was supplemented with blastp alignment to all *Drosophila* polypeptide sequences^52^. Alignment was performed with STARSolo, counting ‘genefulls’, which permits the counting of intronic reads^53^.

Data analysis was performed using Seurat v.5 (Hao et al., 2024). First, all STARSolo-filtered matrices were made into Seurat objects. Empty droplets and non-viable cells with <2000 genes or >20% mitochondrial reads were removed and excluded from subsequent analyses. Cell cycle scores were calculated using the Seurat function CellCycleScoring from the expression of cell cycle markers identified in *D. melanogaster* (https://github.com/hbc/tinyatlas/blob/master/cell_cycle/Drosophila_melanogaster.csv).,

Integration was performed using 3000 genes as anchors to account for variation between source populations (wild *H. melpomene* vs insectary-stock Piano Keys *H. melpomene*), following previously used parameters^31^. Following the validation of merging and integration, the Seurat object was normalised with SCTransform. UMAPs and *k*-means clusters were generated using the first 12 principal components with a resolution of 0.5. Marker genes were selected with FindAllMarkers, using an avg_log2FC cutoff of 0.5. For analysis of the SOP lineage, cells were retained if they expressed known cell type-specific marker genes. The selected cells were then re-normalised and integrated using 3000 genes as anchors similar to the whole dataset.

In order to group scales into “low *ivory*” and “high *ivory”* populations, we normalised *ivory* counts for each sample by mean expression for that sample; this procedure controls for the differences in *ivory* expression levels between samples. We performed pseudobulk differential expression analysis between “high” and “low” populations. GO term enrichment analysis was then performed with PANGEA and ModPhea, looking from predefined categories from levels 10-17^35,54^.

## Supplementary Figures

**Figure S1:**
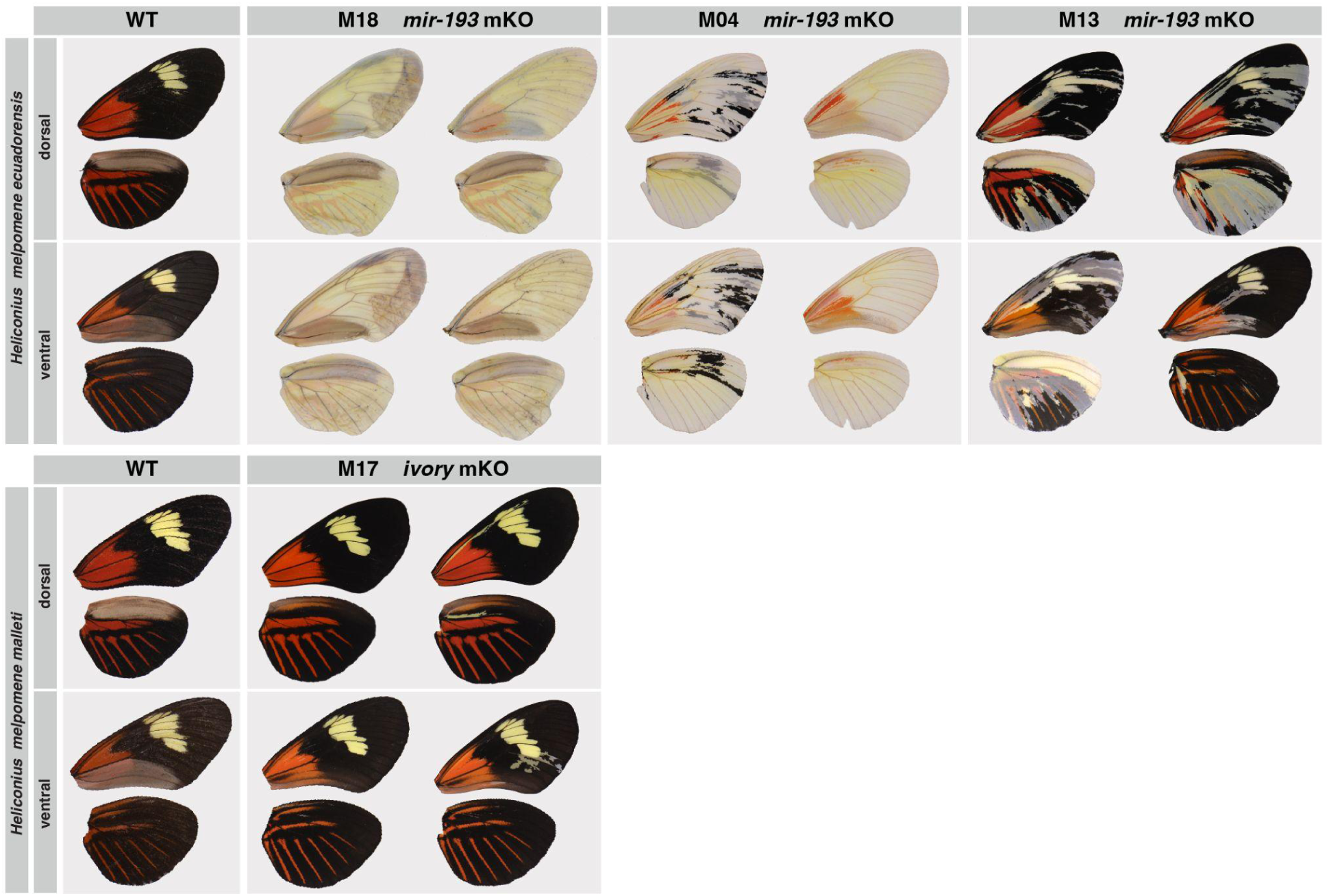
Crispant individuals following somatic mutagenesis of *ivory:mir-193 in H. melpomene ecuadorensis* / *malleti.* These two subspecies have minor differences in the size of the forewing yellow band. Right dorsal wings and left ventral wings were digitally flipped so all wings are shown in the same orientation.

**Figure S2:**
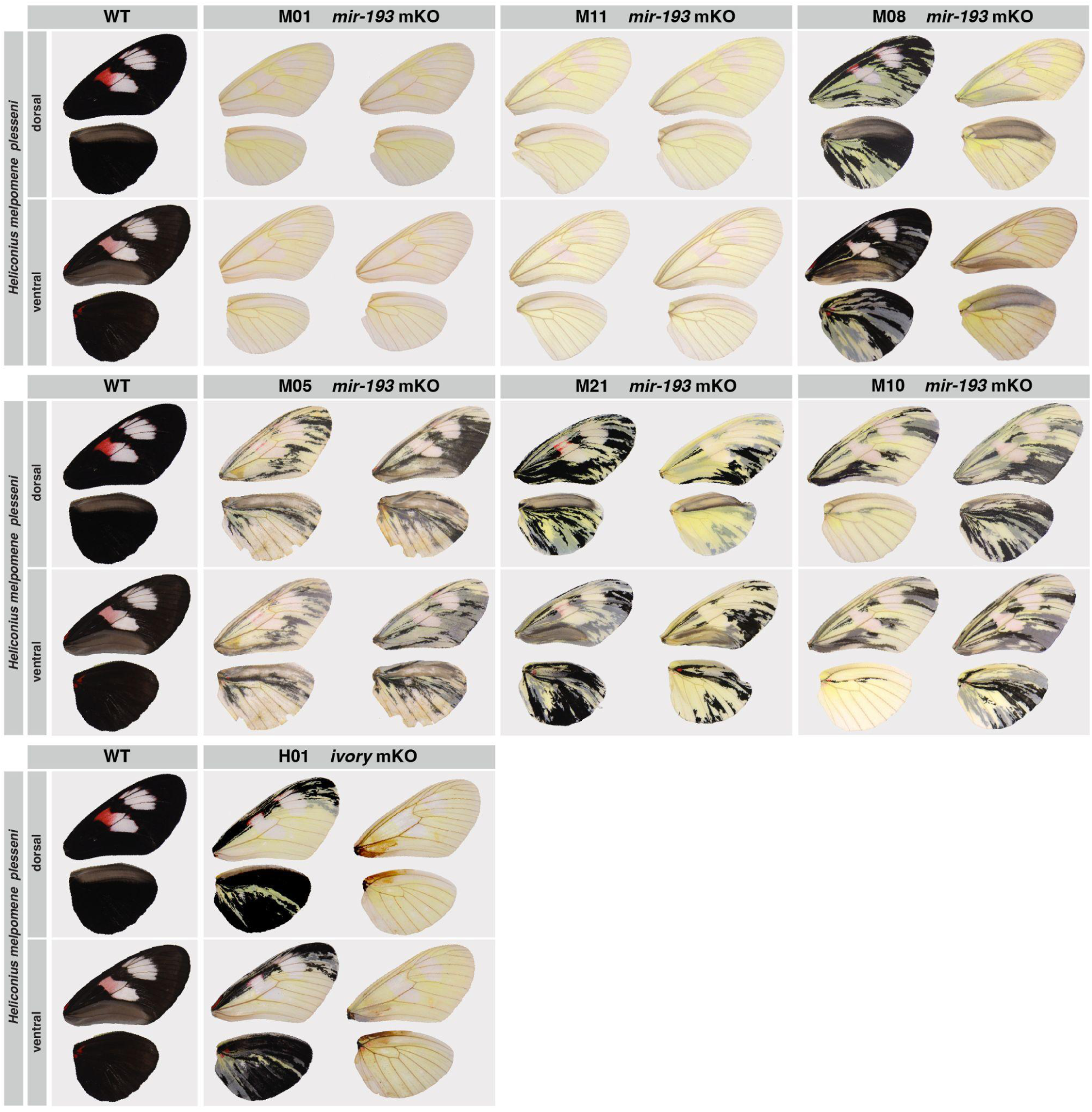
Crispant individuals following somatic mutagenesis of *ivory:mir-193 in H. melpomene plesseni.* Right dorsal wings and left ventral wings were digitally flipped so all wings are shown in the same orientation.

**Figure S3:**
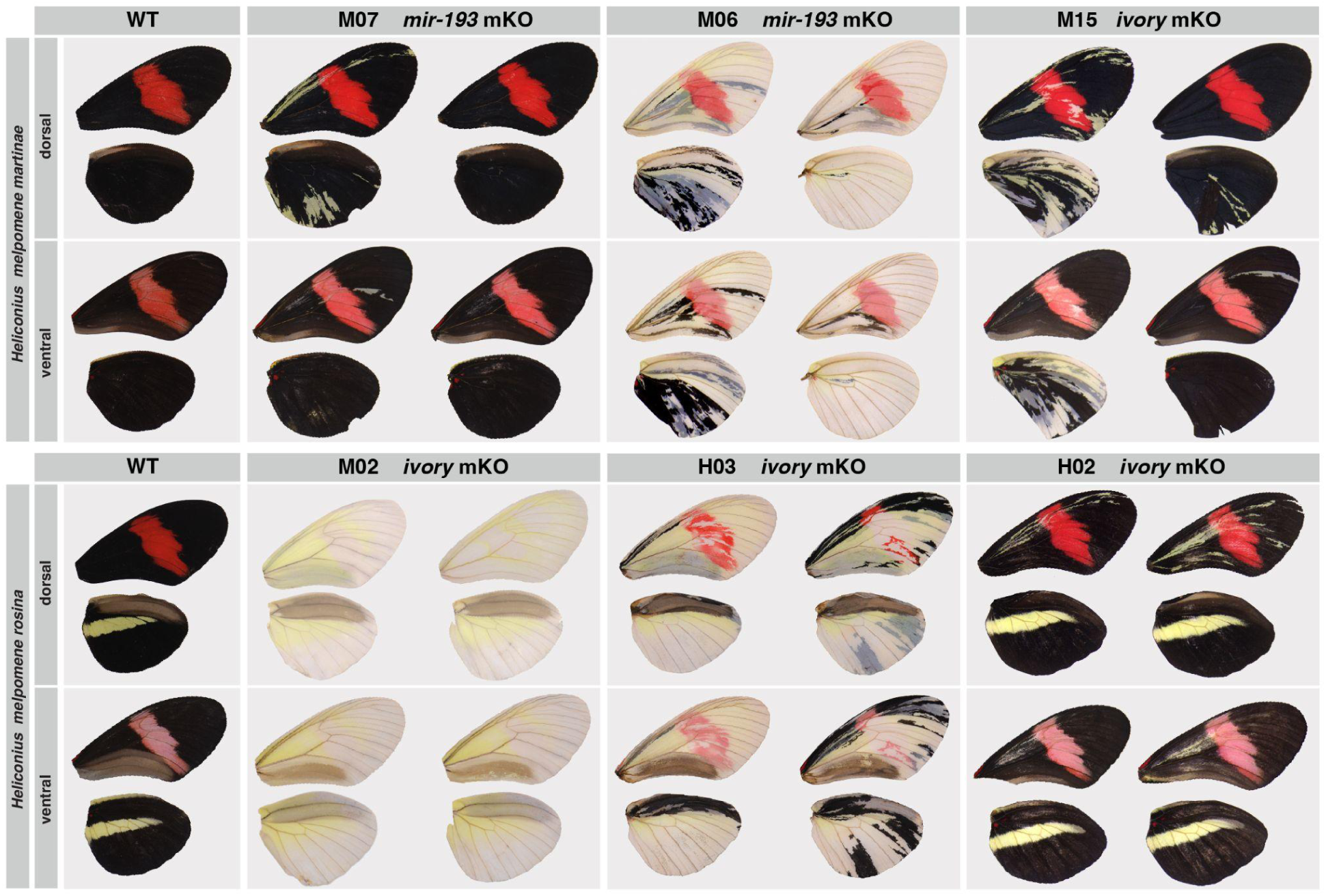
Crispant individuals following somatic mutagenesis of *ivory:mir-193 in H. melpomene martinae* / *rosina.* These two subspecies differ in the presence/absence of a hindwing yellow bar. Right dorsal wings and left ventral wings were digitally flipped so all wings are shown in the same orientation.

**Figure S3:**
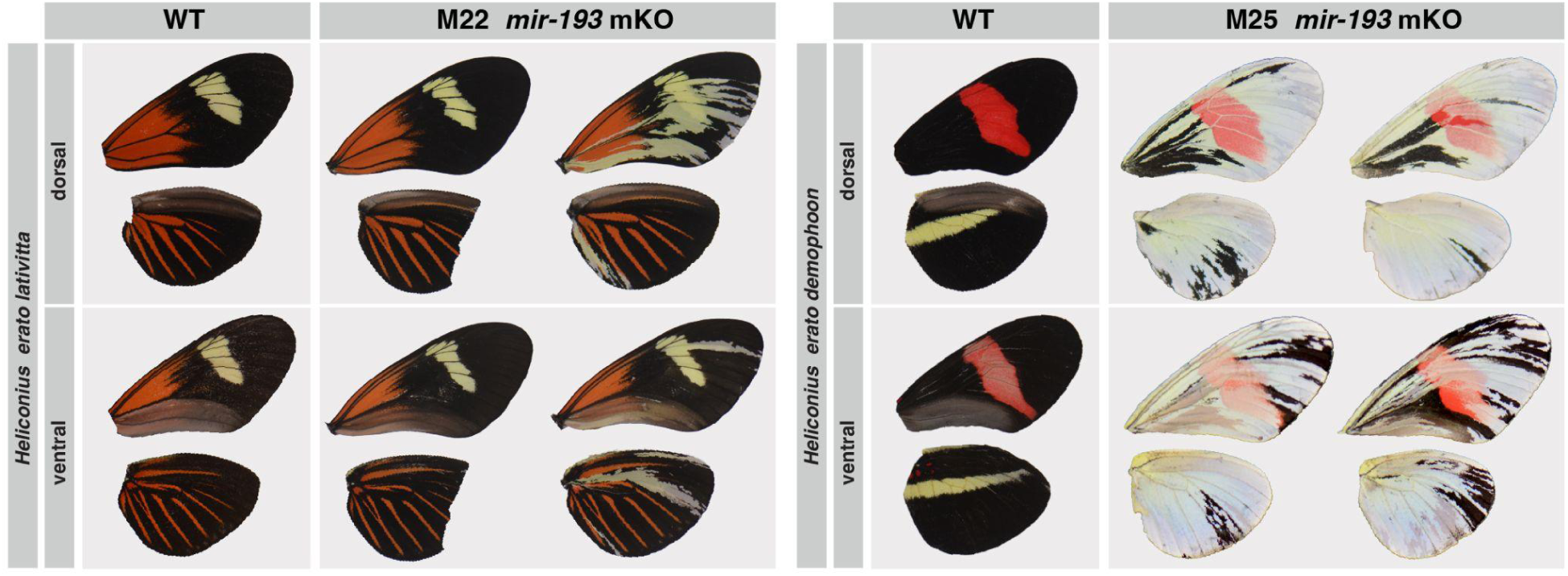
Crispant individuals following somatic mutagenesis of *ivory:mir-193 in the H. erato lativitta* and *H. erato demophoon.* Right dorsal wings and left ventral wings were digitally flipped so all wings are shown in the same orientation.

**Figure S4:**
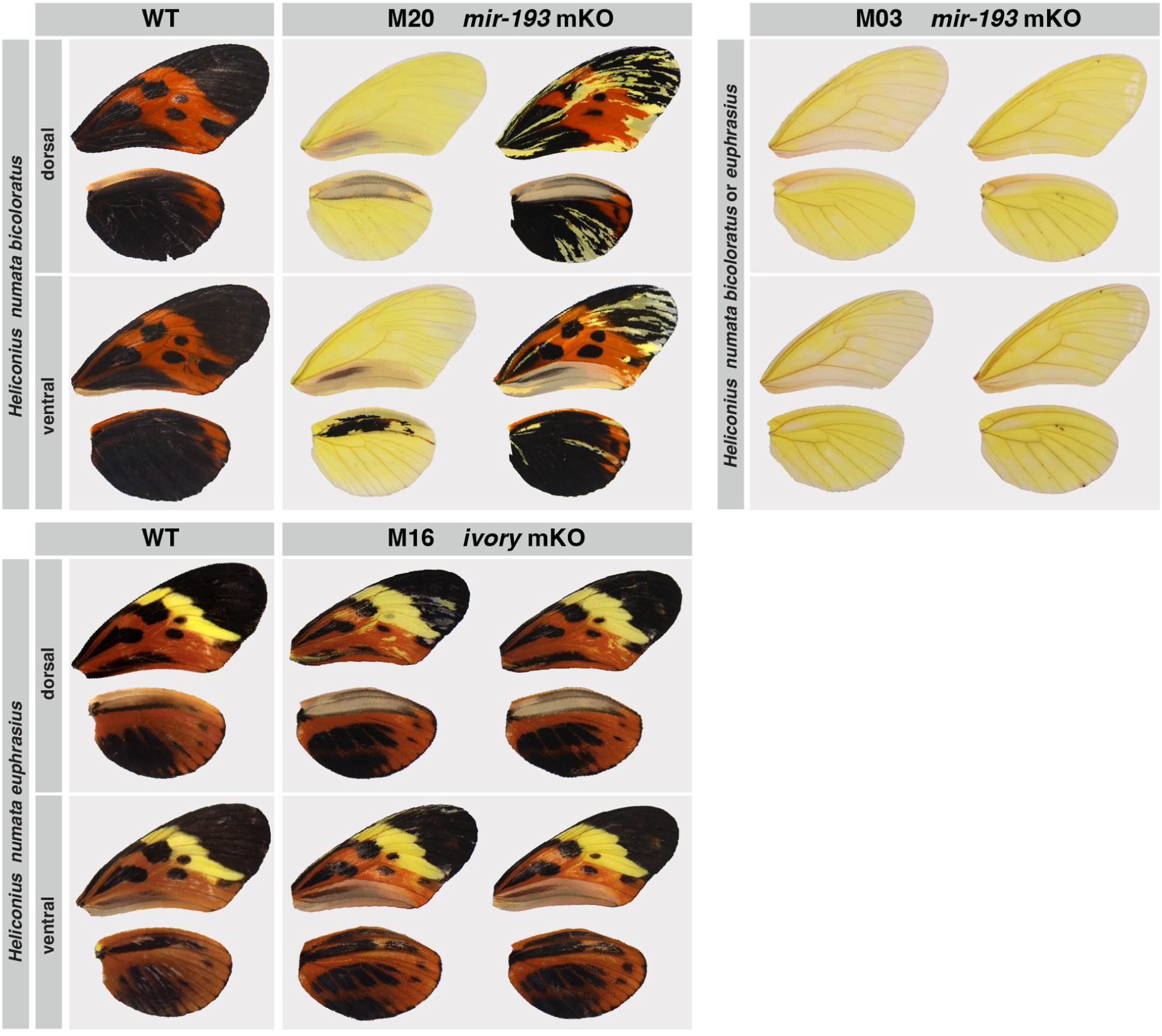
Crispant individuals following somatic mutagenesis of *ivory:mir-193 in H. numata/euphrasius.* The two morphs co-occur in the same colony, and individual M03 can not be assigned to either morph due to its extensive discolouration phenotype. Right dorsal wings and left ventral wings were digitally flipped so all wings are shown in the same orientation.

**Figure S5.**
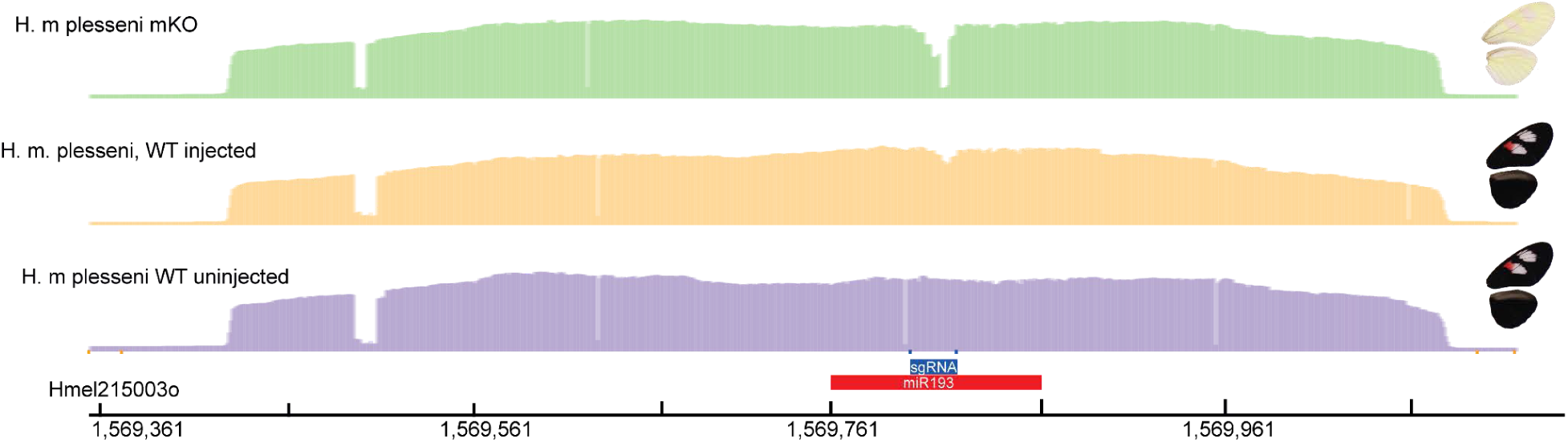
Crispant individuals genotyped by amplicon sequencing. Top row shows read depth of nanopore reads in an extensive crispant. Middle row shows slight loss of coverage at the position of the guide cut site in an individual which was phenotypically wild type. Bottom row shows a wild type individual.

**Figure S6:**
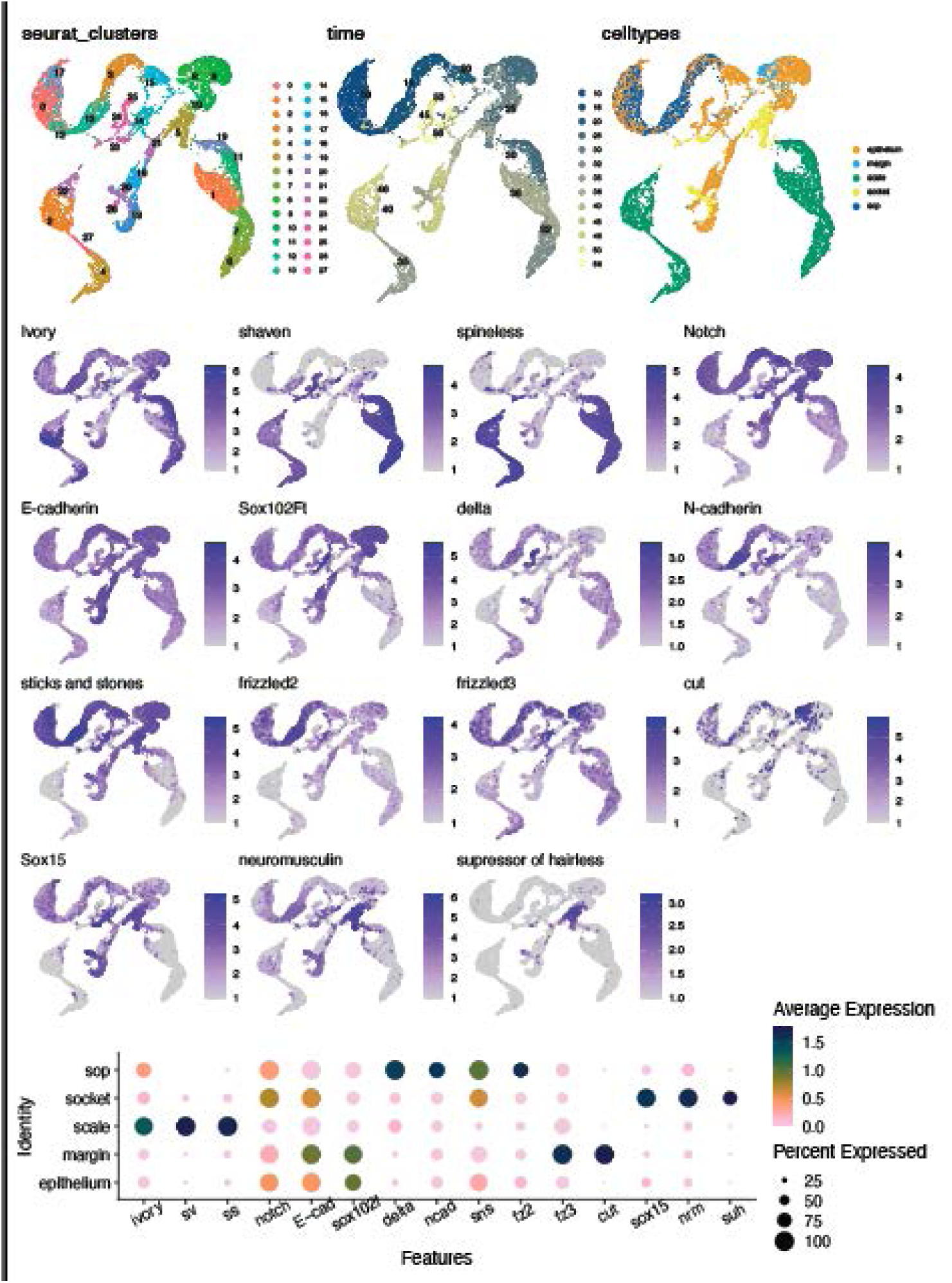
Cell type annotation for integrated data. Top row showing UMAP plots of clusters, time in % pupal development, and assigned cell types based on Loh et al 2025. Second-to-fourth rows show feature plots of multiple celltype markers, and the bottom row shows a dotplot of these markers.

**Figure S7:**
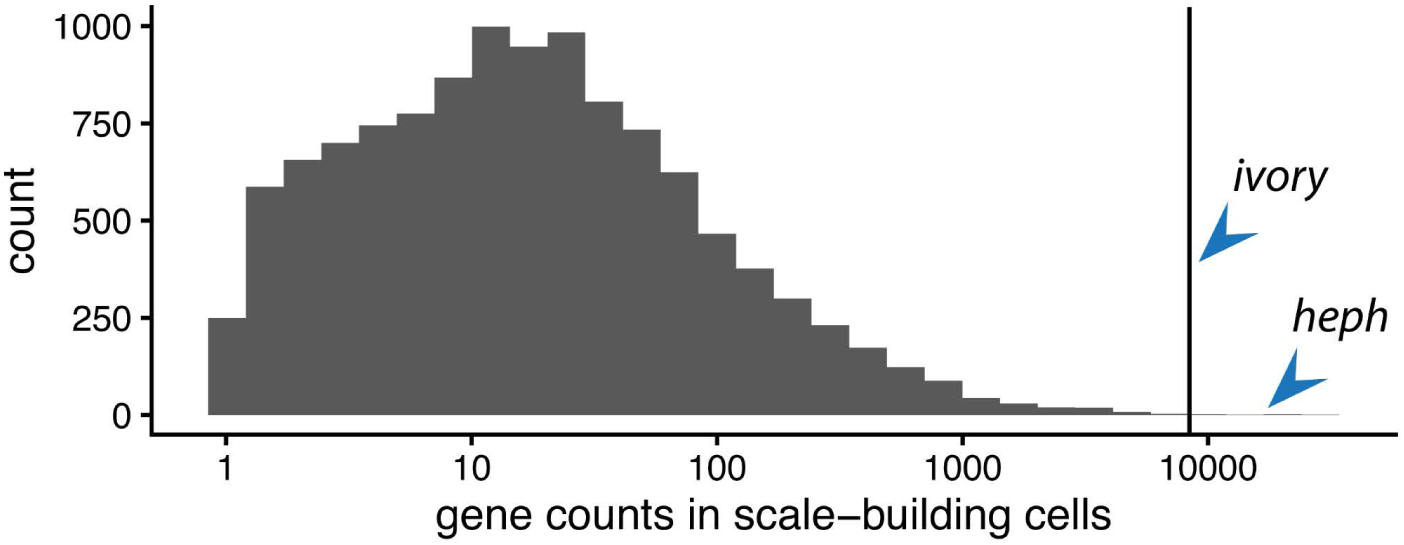
Histogram of gene counts in scale-building cells. Log_10_ of the normalized gene counts on the x-axis, with counts of genes per bin on the y-axis, split across 30 bins. Arrows indicating the top-two genes, *ivory* and *heph*.

**Table S1:**
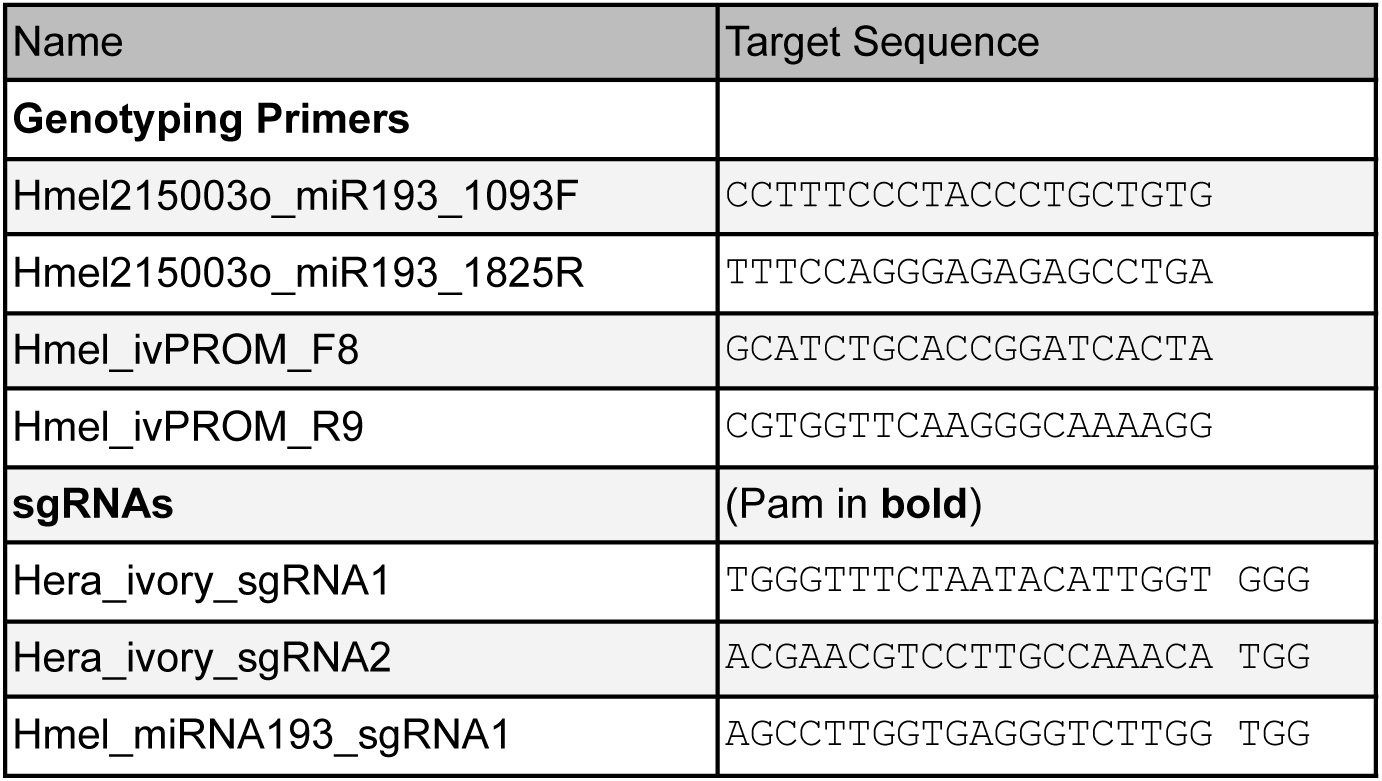
Oligonucleotide sequences.

**Table S2:**
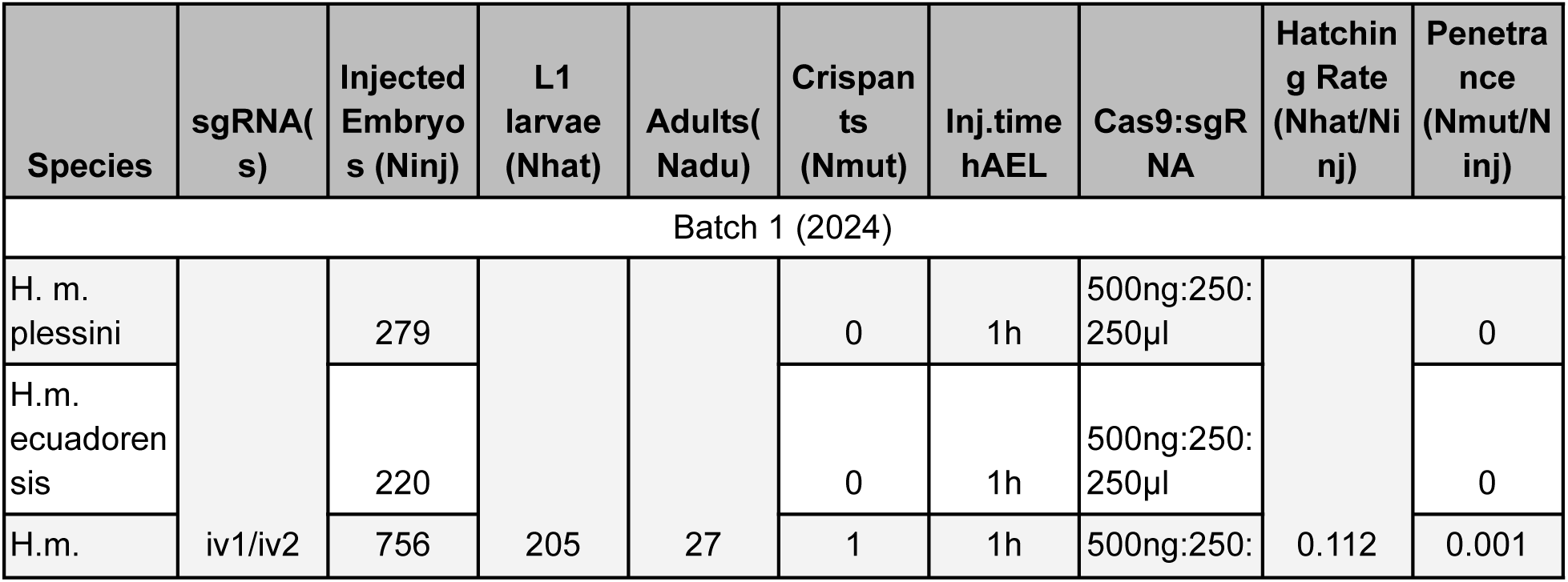
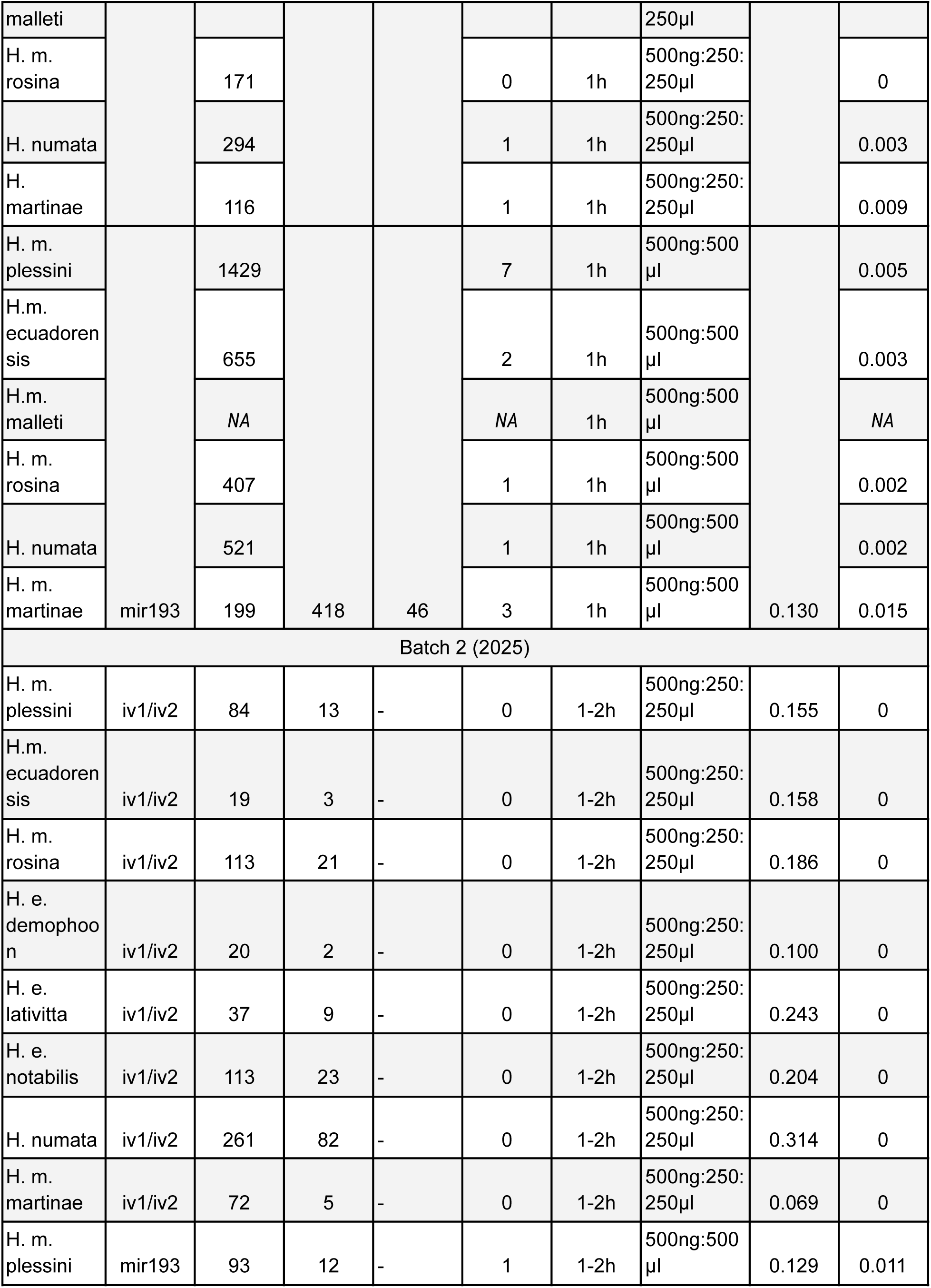
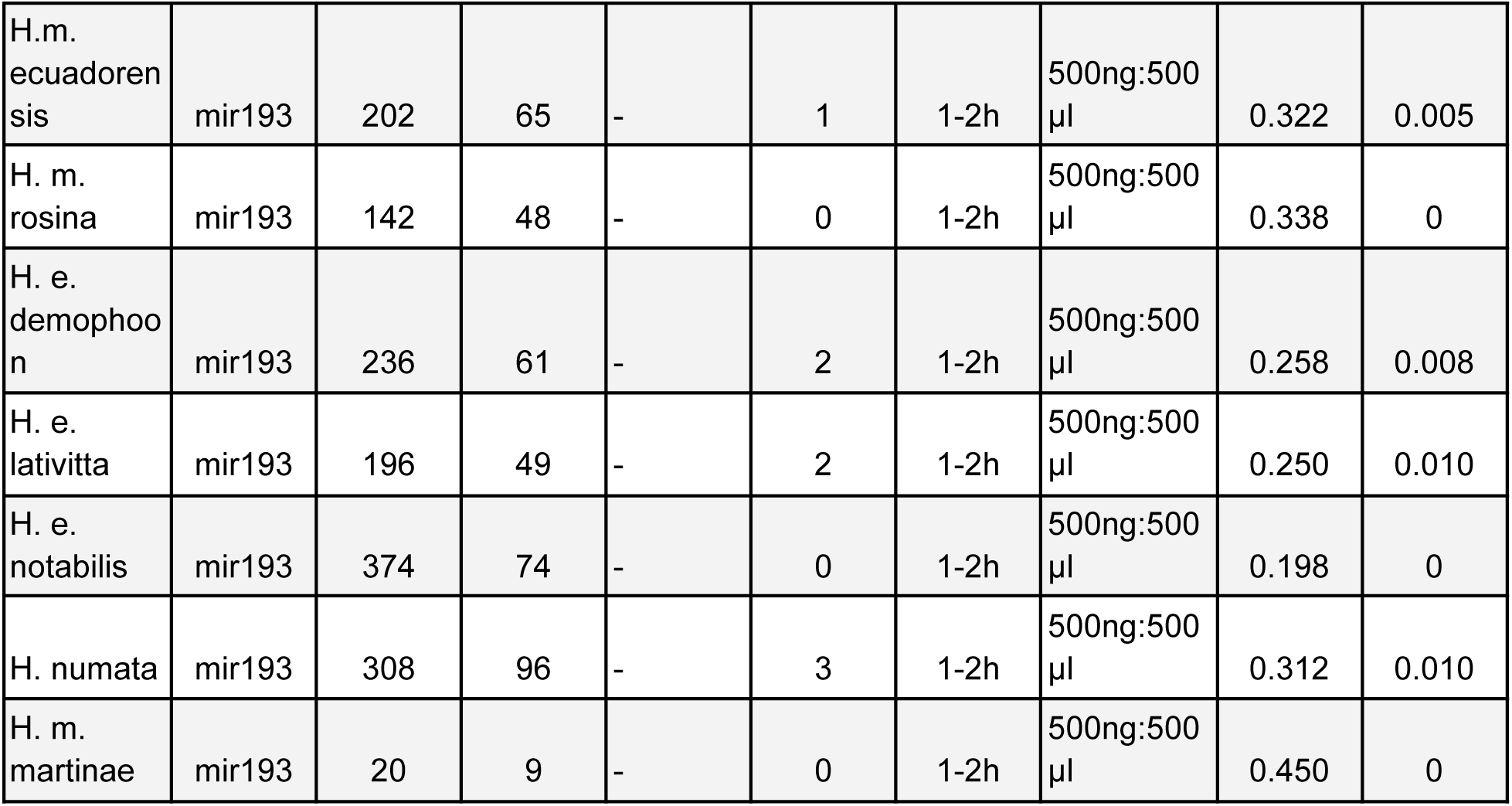
Summary of CRISPR injection experiments related to figure 2, 3 and 4.

**Table S3:**
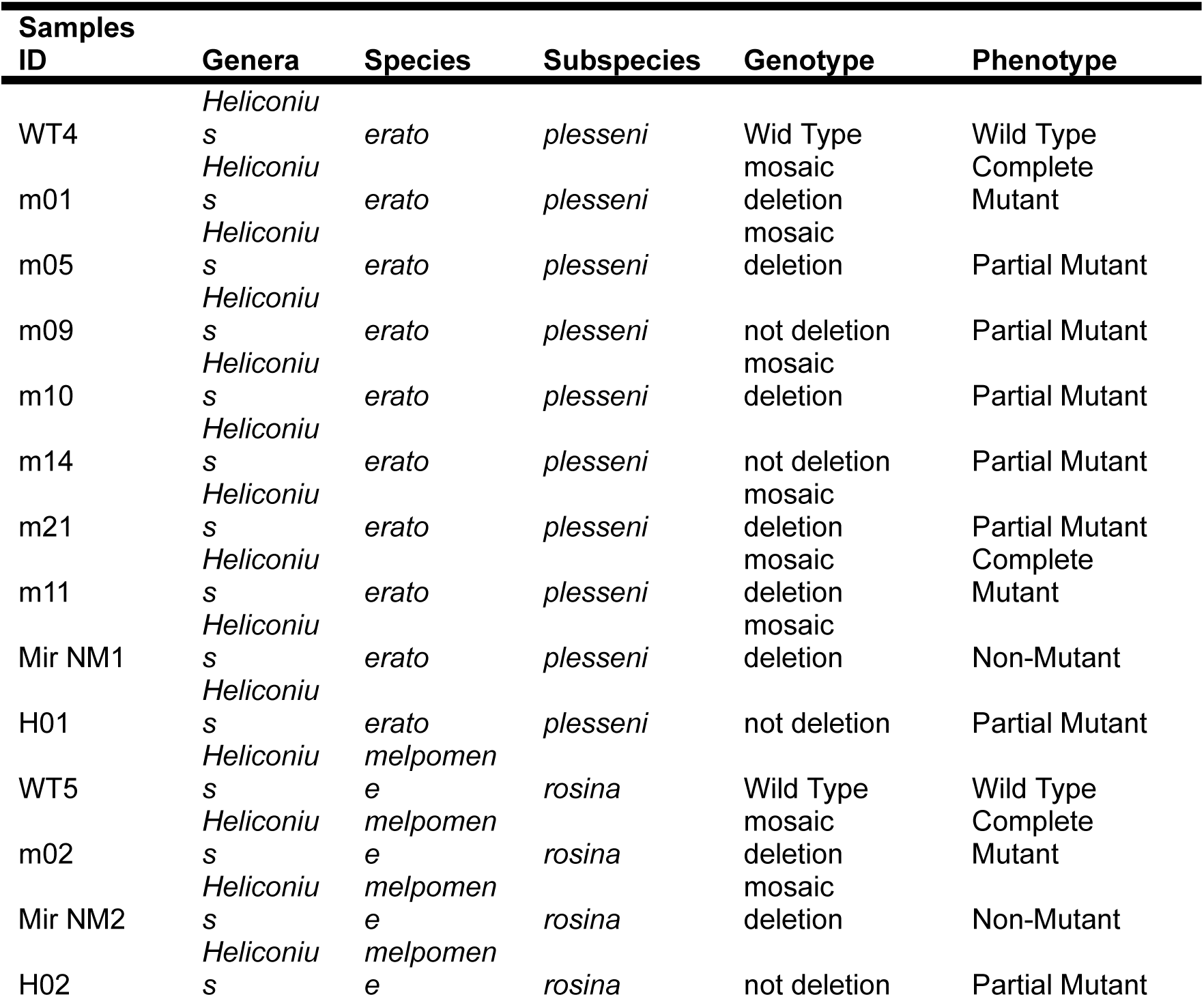
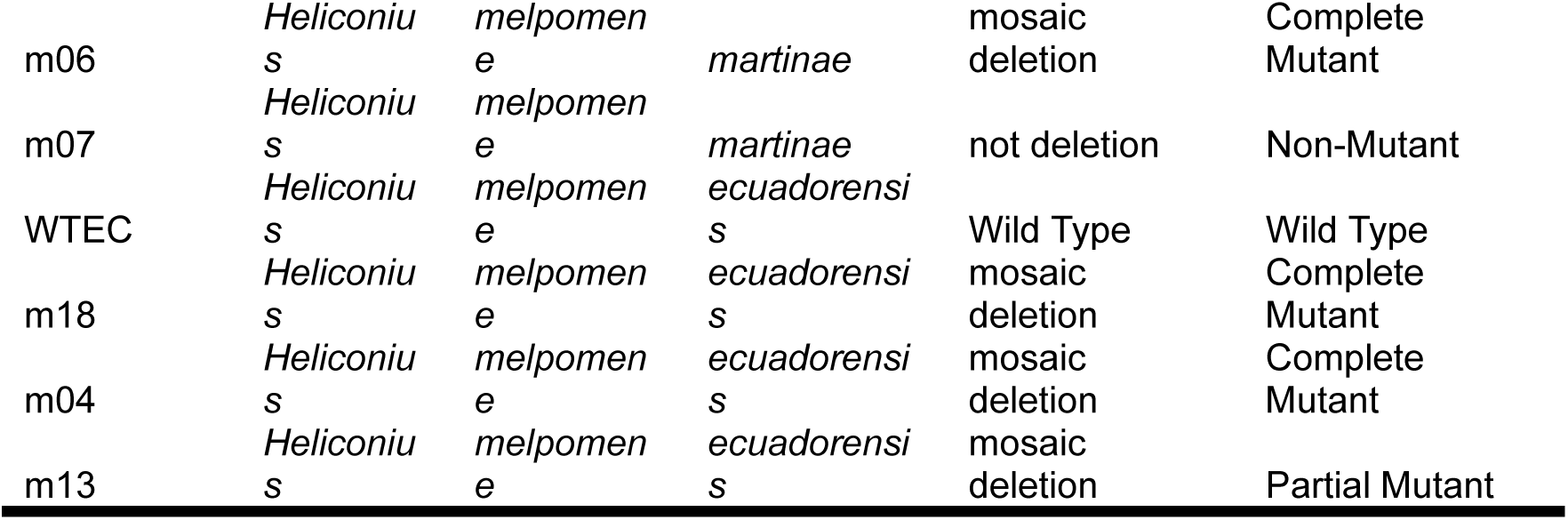
List of wild-type and CRISPR-edited individuals genotyped at the *ivory–mir193* region in *Heliconius erato* and *H. melpomene* populations. The Genotype column indicates whether individuals show a *mosaic deletion* (variable-length deletions surrounding the CRISPR guide cut site) or *no deletion* in the targeted region. The Phenotype column describes the extent of the visible wing-scale mutation: Complete Mutant indicates a fully expressed mutant phenotype in both wings, whereas Partial Mutant indicates asymmetric or incomplete scale transformation. Non-Mutant individuals show no visible phenotypic effect

